# Pleistocene demographic histories dominate contemporary genomic diversity in a continental radiation of Himalayan-Hengduan songbirds

**DOI:** 10.1101/2025.08.28.672032

**Authors:** Yadan Liu, Chih-Ming Hung, Yunrui He, Fei Wu, Dao Yan, Gang Song, Feng Dong

**Author notes:** **Corresponding author:** Feng Dong, Address: No.17 Longxin Road, Kunming.

## Abstract

While population genetic theory expects genetic diversity to scale predictably with population size, mounting empirical evidence has challenged this fundamental prediction. This striking inconsistency has prompted a paradigm shift in our understanding of the determinants of genetic diversity, driving increased efforts to disentangle the relative contributions of Pleistocene demography and linked selection. Here, based on systematic sampling and a unified analytical pipeline, we *de novo* assembled genomes for 120 songbird species breeding in the Himalayas-Hengduan Mountains (HHMs) and conducted population genomic analysis to examine the drivers of their genomic diversity. We observed a 6.5-fold variation in genome-wide heterogeneity and a 16.4-fold variation nucleotide diversity and across species. Notably, these measures of genomic diversity showed no correlation with recent population dynamics, current population size, or other contemporary factors—such as natural selection, elevational distribution, or life-history traits. Instead, historical demography strongly predicted genetic diversity, with ancestral population size during the late Pleistocene emerging as the sole correlate: larger ancestral sizes consistently coincided with higher diversity. These findings underscore the critical influence of historical demography on contemporary genetic diversity in natural populations—an insight essential for designing effective conservation strategies.

## Introduction

Global biodiversity is currently confronting an unprecedented crisis, characterized by species extinction rates exceeding 100 times those observed in pre-human eras (1), coupled with widespread population declines across extant species (2). Compounding this crisis, genetic diversity—a fundamental component of biodiversity—has declined by approximately 6% to 10% over the past century (3,4), with long-term projections suggesting potential losses exceeding 50% (5). Such declines directly threaten population viability, species’ evolutionary potential, and ecosystem resilience (6), while undermining the United Nations’ post-2020 genetic conservation targets (7). This alarming scenario necessitates a deeper understanding of the determinants governing genetic diversity, which is critical for both elucidating underlying mechanisms of biodiversity loss and developing effective conservation strategies.

The neutral theory of molecular evolution predicts that genetic diversity scales with population abundance (8), yet empirical studies consistently reveal deviations from this expectation (9, 10). Current theoretical frameworks center on two competing paradigms: Pleistocene demography versus natural selection (11, 12). Climates in the late Pleistocene were marked by high-amplitude glacial–interglacial cycles (13), which likely triggered severe demographic fluctuations—particularly population bottlenecks. Such bottlenecks consistently lead to prolonged reductions in genetic diversity, owing to the extended recovery time required after population decline (14). Alternatively, linked selection could reduce diversity at neutral sites through genetic linkage, driving genome-wide diversity below neutral expectations (11). Beyond these dominant mechanisms, species-specific traits (e.g., reproductive strategies) can modulate diversity by influencing ecological resilience (10), while environmental factors (e.g., elevation) may further contribute to genomic variation (15). Despite these insights, the relative contributions of these processes remain uncertain, as inconsistent data quality and the challenge of disentangling ecological and evolutionary confounders persist (9).

Songbirds (Passeriformes, Aves), representing over 45% of extant avian diversity, constitute a globally significant radiation and conservation priority (16, 17). The Himalayas–Hengduan Mountains (HHMs), a global biodiversity hotspot, provides an ideal study system with its exceptional songbird diversity and extreme habitat heterogeneity (18). This mountain system’s dramatic elevational gradients and unparalleled bioclimatic variation have imposed strong selective pressures throughout songbird evolutionary history (19), establishing the HHMs as a natural laboratory for investigating determinants of genetic diversity.

This study employs a comprehensive framework that integrates systematic sampling and high-resolution genomic analyses across 120 breeding songbird populations in the HHMs, aiming to elucidate the key drivers of genetic diversity. Our approach specifically evaluates the relative contributions of Pleistocene demographic history versus selective processes in shaping genomic variation. These competing hypotheses yield distinct, testable predictions: the demographic model posits that genetic diversity correlates with Pleistocene minimal population sizes, whereas the selection model proposes that diversity is associated with the efficacy of natural selection (11, 20).

## Results

### Genome assembly and population genomic dataset

To quantify genetic diversity, we first performed *de novo* assembly of a reference genome for each of the 120 studied bird species. The resulting genomes exhibited high quality, with average sequencing coverage ranging from 33.1× to 72.4×, scaffold N50 values between 70.9 and 106.4 Mbp, and strong completeness—92.5% of assemblies achieved BUSCO (Benchmarking Universal Single-Copy Orthologs) completeness scores of ≥ 90% (*SI Appendix*, Table S1). Leveraging these high-quality genomic resources, we constructed a population genomic dataset by mapping resequencing data—obtained from five individuals per species at coverages ranging from 5.6× to 30.9×—to their corresponding reference assemblies (*SI Appendix*, Table S2).

To ensure analytical consistency, we implemented a rigorous data normalization protocol in which the original high-coverage sequencing data (used for genome assembly) were systematically downsampled to 30% of their original depth prior to population genomic analyses. This approach effectively mitigated potential biases arising from highly heterogeneous sequencing coverage.

### Sampling design and diversity metrics

We addressed sampling bias in genomic diversity calculations through two key strategies, specifically minimizing potential confounding effects of spatial genetic structure—particularly isolation-by-distance patterns (15), a common challenge in genetic diversity studies. First, for all but one species, we implemented a localized sampling approach for all but one species, collecting five individuals per species exclusively from either the eastern Himalayas or the southern Hengduan Mountains (*SI Appendix*, Fig. S1 and Table S2), avoiding cross-regional sampling. Second, we characterized genomic diversity using two metrics: population-level nucleotide diversity (*π*) and individual-based heterozygosity (*h*), both of which perform well under spatial sampling (21).

### Interspecific variation in genetic diversity

Genomic analyses across 120 species revealed remarkable interspecific variation in genome-wide diversity (*π*: 16.4-fold; *h*: 6.5-fold; Fig. 1). Notably, this variation lacked phylogenetic signal or taxonomic clustering (*π*: Blomberg’s K = 0.26, permutation test *p* = 0.860; *h*: Blomberg’s K = 0.29, permutation test *p* = 0.715). The extremes of this spectrum were, for instance, represented by the yellow-eyed babbler (*Chrysomma sinense*; *π* = 2.32×10^-3^) and blue-winged minla (*Actinodura cyanouroptera*; *π* = 3.81×10^-2^), which showed the lowest and highest diversity values, respectively.

**Fig. 1.**
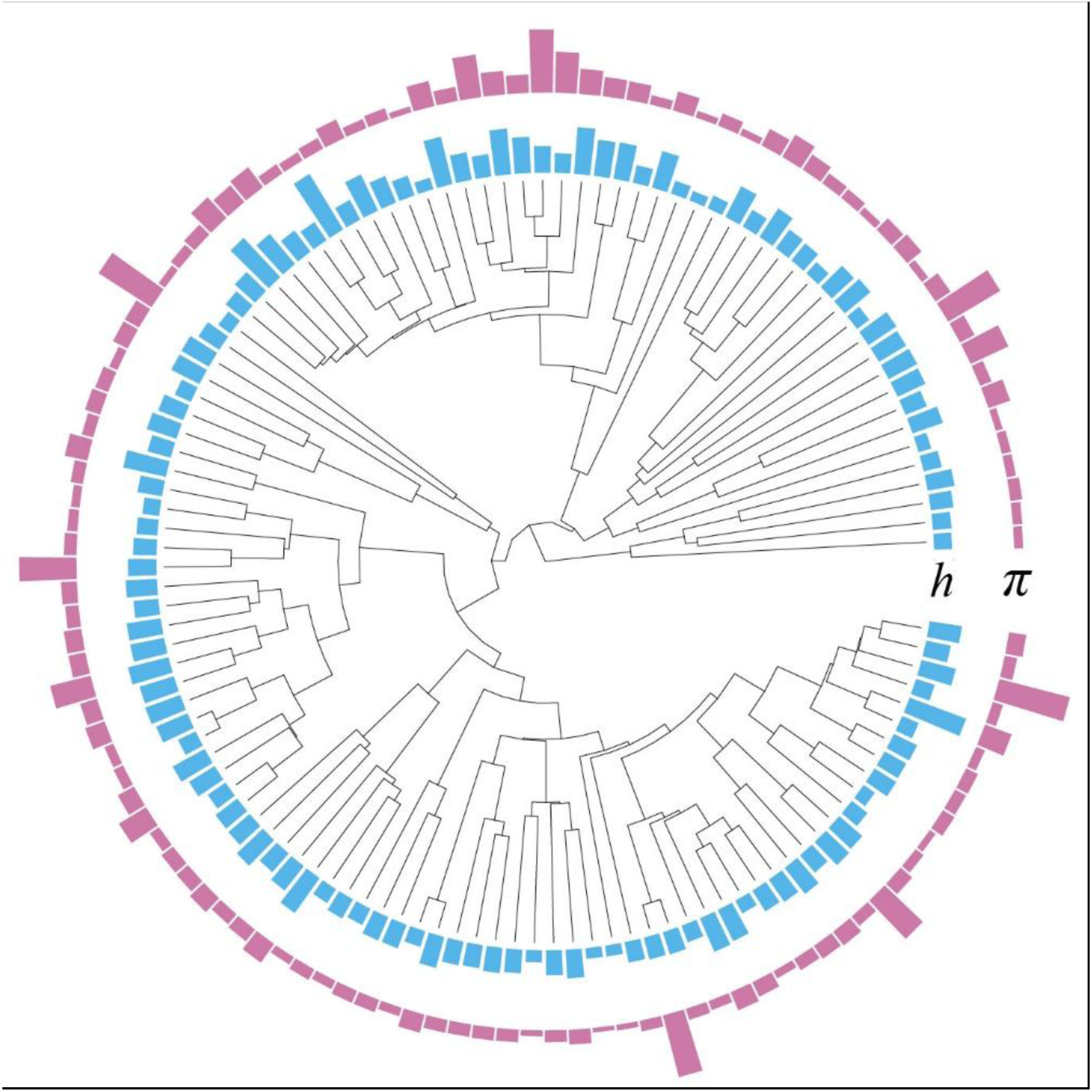
Genome-wide nucleotide diversity (*π*) and heterogeneity (*h*) across 120 songbird species. Phylogeny was constructed using the Jetz backbone (72) from www.birdtree.org. Observed variation spans 16.4-fold for *π* and 6.5-fold for *h* across species.

### Pleistocene demography as the primary driver of diversity

To elucidate the drivers underlying the observed patterns of genetic diversity, we implemented a Bayesian phylogenetic generalized linear mixed model framework (22). We identified ancestral population size as the primary predictor of genetic diversity across HHM songbirds. For each species, we estimated ancestral population size as the harmonic mean of effective population sizes (*N_e_*) over the late Pleistocene using a linkage-based hidden Markov model (pairwise sequentially Markovian coalescent, PSMC) (23). Our analysis revealed strong positive correlations between the historical harmonic mean of *N_e_* and both *π* (*z* = 0.34 [95% credible interval (CI): 0.14–0.54], *pMCMC* < 0.001) and *h* (*z* = 0.44 [0.26–0.64], *pMCMC* < 0.001; Fig. 2 and 3, and *SI Appendix*, Fig. S2 and S3).

**Fig. 2.**
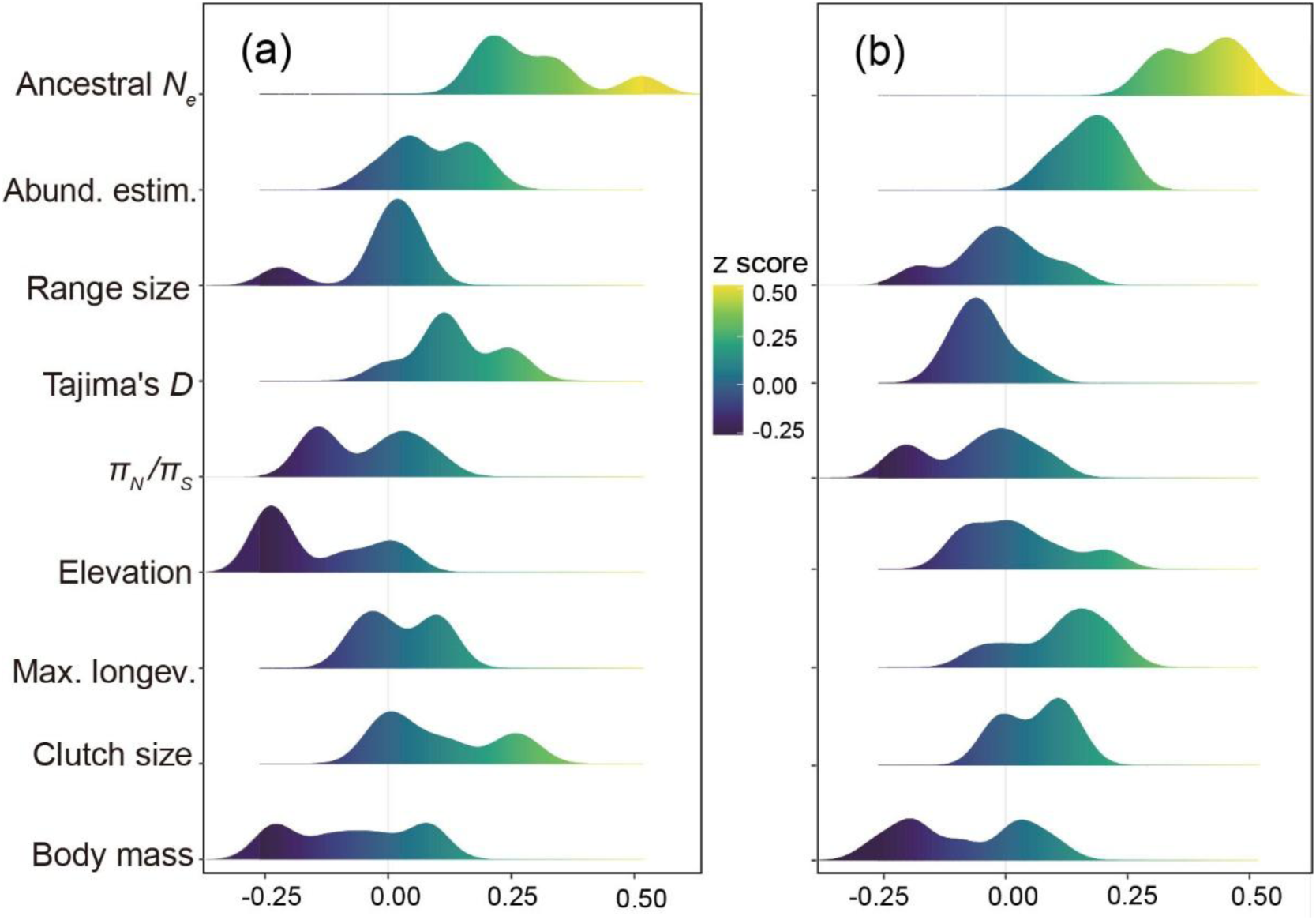
Predictors and their effects on genomic diversity in the 120 studied songbirds. Distribution of the estimated coefficients for the potential explanatory variables of genome-wide nucleotide diversity (a) and heterozygosity (b) across the 120 studied birds. These variables include long-term *N_e_*, population abundance estimate (Pop. abund.), range size, Tajima’s *D*, natural selection (represented by *π*_N_/*π*_S_), elevation, and life-history traits (maximum longevity, Max. longev.; clutch size; and body mass). The results suggest that long-term *N_e_* is the only significant predictors of both metrics of genetic diversity.

**Fig. 3.**
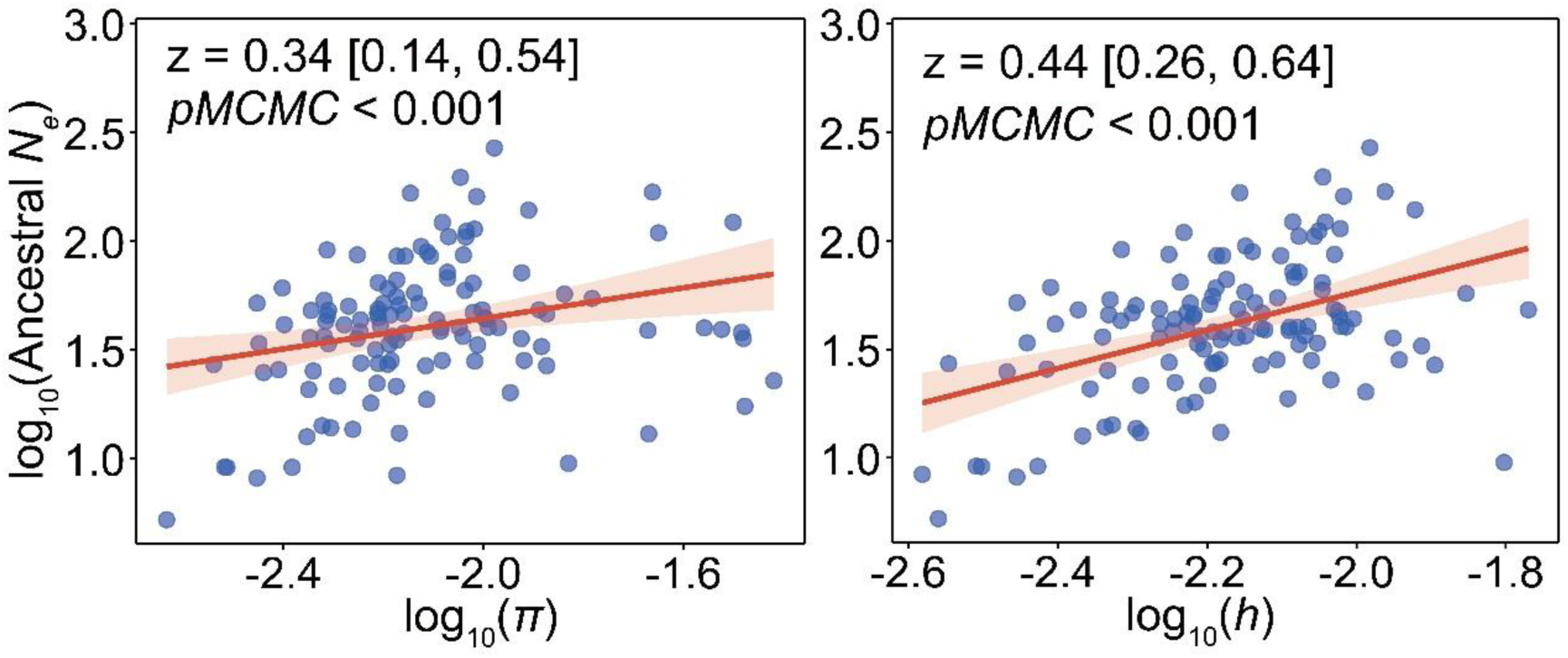
Relationship between long-term effective population size (*N_e_*) and genetic diversity across 120 songbird species. Log-transformed long-term *N_e_* correlates with log-transformed genome-wide nucleotide diversity (*π*) and heterozygosity (*h*). Bayesian phylogenetic generalized linear mixed models were used to compute *z*- scores with 95% confidence intervals (shaded pink) and assess statistical significance. The red line indicates the model fit.

### Limited effects of contemporary population size and recent expansions

To assess the influence of contemporary population size, we evaluated two surrogate measures: estimated abundance and geographic range size. Regarding abundance estimate, neither *π* nor *h* showed a significant relationship (*π*: *z* = 0.15 [- 0.10, 0.39], *pMCMC* = 0.254; *h*: *z* = 0.22 [-0.02, 0.48], *pMCMC* = 0.076; Fig. 2 and 3, and *SI Appendix*, Fig. S2 and S3). Similarly, for range size, neither measure of genetic diversity was significantly associated (*π*: *z* = -0.08 [-0.33, 0.16], *pMCMC* = 0.480; *h*: *z* = -0.04 [-0.25, 0.22], *pMCMC* = 0.740; Fig. 2 and 3, and *SI Appendix*, Fig. S2 and S3).

Recent demographic trends, as inferred from Tajima’s *D* values, also showed no significant relationship (*π*: *z* = 0.15[-0.04, 0.33], *pMCMC* = 0.104; *h*: *z* = -0.02 [-0.22, 0.12], *pMCMC* = 0.796; Fig. 2 and 3, and *SI Appendix*, Fig. S2 and S3). Notably, 95% of species (114/120) exhibited negative Tajima’s *D* values *(SI Appendix*, Table S1 and Fig. S4), with the magnitude of these values suggesting post-bottleneck expansions.

### Minimal influence of natural selection and elevational distribution

To examine the role of natural selection in determining genetic diversity, we calculated the ratio of nonsynonymous to synonymous nucleotide diversity (*π*_N_/*π*_S_) in coding regions for each bird as a measure of selection efficacy (24). This ratio reflects the strength of purifying selection: efficient purifying selection reduces *π*N/*π*S, while weaker selection results in higher values. Our analyses revealed no significant correlation between *π*_N_/*π*_S_ ratios and genetic diversity (*π*: *z* = -0.02 [-0.19, 0.15], *pMCMC* = 0.854; *h*: *z* = -0.01 [-0.18, 0.16], *pMCMC* = 0.970; Fig. 2 and *SI Appendix*, Fig. S2 and S3). Similarly, elevational distribution proved to be a poor predictor of diversity (*π*: *z* = -0.20 [-0.41, 0.01], *pMCMC* = 0.076; *h*: *z* = -0.03 [-0.22, 0.16], *pMCMC* = 0.774; Figs. 2 and *SI Appendix*, Fig. S2 and S3).

### Nonsignificant role of life-history traits

We found no significant relationship between life-history traits and genetic diversity in HHM songbirds (Fig. 2 and *SI Appendix*, Fig. S2 and S3). Neither clutch size (*π*: *z* = 0.12 [-0.06, 0.33], *pMCMC* = 0.222; *h*: *z* = 0.06 [-0.15, 0.23], *pMCMC* = 0.572), maximum longevity (*π*: *z* = -0.03 [-0.22, 0.17], *pMCMC* = 0.762; *h*: *z* = 0.08 [- 0.11, 0.29], *pMCMC* = 0.398), nor body mass (*π*: *z* = -0.06 [-0.26, 0.12], *pMCMC* = 0.588; *h*: *z* = -0.06 [-0.25, 0.11], *pMCMC* = 0.522) showed significant associations with genetic diversity measures.

## Discussion

Our study leverages population genomic data from 120 HHM songbirds to assess the determinants of genetic diversity, with a specific focus on the relative roles of historical demography and natural selection. We demonstrate that patterns of genetic diversity in these birds are predominantly shaped by demography during the late Pleistocene—rather than by contemporary ecological or genetic factors—supporting the classical hypothesis that Pleistocene climatic fluctuations fundamentally influenced present-day diversity (25).

During the cyclical glacial periods of the late Pleistocene (26), harsh environmental conditions likely reduced HHM songbird populations to minimal sizes (27, 28), leaving a lasting imprint on their genetic diversity (11). Consistent with this, we found strong positive correlations between the harmonic mean of *N_e_* estimates during the late Pleistocene and both metrics of genetic diversity (*π* and *h*), indicating that species with larger Pleistocene *N_e_* retained higher diversity. This parallels findings from other empirical studies (29–32).

Given the dominant influence of Pleistocene demography, we expected recent demographic expansions and current population size to have minimal effects. This is because rapid increases in census population size contrast sharply with the slow recovery of genetic diversity (11). Populations recovering from severe bottlenecks require extended evolutionary timescales—often millions of years—to restore mutation-drift equilibrium (14). Our results support this expectation: neither contemporary population size (measured via abundance estimate or range size) nor recent expansions (inferred from Tajima’s *D*) significantly predicted diversity. A similar pattern is evident in *Homo sapiens*—despite a dramatic expansion from a few thousand to 7.9 billion individuals, long-term *N_e_* remains low (∼25,000) (12). These findings are consistent with broader taxonomic trends (32–34), underscoring the lasting impact of long-term demographic history, rather than recent changes, on genetic diversity (30, 35).

The contribution of natural selection to genetic diversity patterns remains debated (11). Although Corbett-Detig et al. (20) identified linked selection as a key driver of eukaryotic diversity, subsequent analyses could not confirm this effect (36). Our study, using *π*_N_/*π*_S_ as a selection proxy, found minimal evidence for linked selection shaping genome-wide diversity in HHM songbirds - despite their broad ecological range spanning tropical to near-boreal elevations (225-4,250 m; *SI Appendix*, Table S1 and Fig. S5). This contrasts with findings in butterflies (37), suggesting taxon-specific variation in selection’s influence that warrants broader phylogenetic investigation.

Our analyses demonstrate that elevational distribution poorly predicts genetic diversity in HHM songbirds, challenging assumptions in macrogenetics - a field studying landscape-scale diversity patterns (38). Current macrogenetic studies show inconsistent diversity gradients along environmental clines (15, 39), potentially reflecting methodological limitations of common markers: mitochondrial DNA (vulnerable to maternal inheritance and selection biases) (40) and microsatellites (with variable mutation rates compromising cross-species comparisons) (11). By analyzing genome-wide data from 120 species, we provide robust evidence that elevation minimally influences diversity patterns in these birds, underscoring the value of genomic approaches for macroecological insights.

Although life-history traits (especially reproductive strategies) can shape genetic diversity by affecting environmental responses (10), this fails to explain variation among species with similar strategies (37). Songbirds present a clear case, as they uniformly show low fecundity and high parental care (41). Accordingly, we detected no significant association between life-history traits and genetic diversity in HHM songbirds.

In summary, our study establishes historical demography - not contemporary ecological or genetic factors - as the primary driver of genetic diversity in HHM songbirds. This supports the classical view that past demographic events, especially those triggered by environmental changes, fundamentally shape genetic diversity (25, 42). Consistent with recent work (43), we find contemporary factors like population size, life-history traits, elevation, and selection efficiency poorly predict diversity patterns. These results carry crucial conservation implications: populations maintain evolutionary potential through historical genetic legacies despite mounting anthropogenic threats (44), emphasizing the need to integrate demographic history into conservation planning and genetic assessments of population viability.

## Materials and Methods

### Genomic sequencing and diversity estimation

To obtain reference genomes, we sampled one individual for each bird species from the HHM region and generated genomic sequences via paired-end sequencing with an inset size of 350 bp and a read length of 151 bp on a BGI DNBSEQ platform. We cut adapters with Cutadapter v2.10 (45) and filtered low-quality reads (e.g., Phred score < 15) with Seqtk (46). We aligned quality-controlled reads into contiguous sequences (contigs) via Abyss 2.0 (47). To increase genomic continuity, we assembled these contigs into pseudochromosomes via synteny with the zebra finch assembly (GCA_008822105.2) in Satsuma v2.0 (48). We searched for passerine orthologs (passeriformes_odb10) via BUSCO v5.7.1 (49) to assess genomic completeness.

We compiled a population genomic dataset consisting of five genomes per species, sampled from local communities either in the eastern Himalayas (e.g., the Nanga Bawa Mountain) or the southern Hengduan Mountains (e.g., the Gaoligong Mountain), but not from both regions. The median spatial sampling interval within each species ranged from 0 to 342 km, with larger intervals primarily occurring in the geographically elongated Gaoligong Mountain. For individuals sampled from the same community, sequences originally intended for genome assembly were included in the final population genomic dataset after downsampling via Seqtk to minimize potential biases from exceptionally high genomic coverage. Following the same quality control steps applied in the reference genome assembly process—including adapter trimming and low-quality base removal—we performed read mapping, sorting, and deduplication against species-specific reference genomes. These steps were carried out using Burrows–Wheeler Aligner v0.7.17 (50), Sambamba v0.6.6 (51), and the Genome Analysis Toolkit (GATK) v4.0.3 (52).

We estimated genome-wide π using ANGSD v0.928 (53) with a 50-kb nonoverlapping window size for each species, based on the obtained binary alignment map (BAM) files. The analysis was performed with the following parameters: ‘-GL 1 - doSaf 1 -remove_bads -only_proper_pairs 1 -minMapQ 30 -minQ 20 -minInd 5’, followed by ‘realSFS’ and ‘thetaStat’ for downstream calculations. Individual heterozygosity (*h*) was computed following the standard ANGSD workflow (http://www.popgen.dk/angsd/index.php/Heterozygosity). Population-level *h* was then derived by averaging values across five individuals per species.

### Demographic variables

To investigate the relationship between ancestral population size and genetic diversity, we estimated the harmonic mean of historical *N_e_* values during the late Pleistocene (20-500 Kya) using the PSMC method (23). For each species, we analyzed high-coverage autosomal de novo assemblies with parameter settings revised from Nadachowska-Brzyska et al. (27) (N30–t5–r5–p “1+1+1+1+30∗2+4+6+10”). These parameters were selected as an optimal strategy to avoid artifacts such as large *N_e_* peaks followed by apparent population collapses (54). Prior to PSMC analysis, we prepared input files using BCFtools v0.1.16 (55) with the “-C50” option. Following PSMC documentation recommendations, we applied minimum and maximum read depth filters set to one-third and twice the average genome coverage, respectively.

For recent population dynamics, we calculated Tajima’s *D* statistics using 50 kb sliding windows with consistent parameters (as in our genome-wide estimation) through the ANGSD program. Tajima’s *D* values near zero suggest stable population size, while significantly positive or negative values indicate population contraction or expansion, respectively.

Modern population size was accessed using two proxies: (1) abundance estimates from (56), though we acknowledge potential unquantifiable biases (57); and (2) range size (58), which is a well-established positive correlate of census population size (59).

### Efficiency of purifying selection

To investigate how linked selection influences genetic diversity variation in the studied songbirds, we calculated the πN/πS ratio in the coding region for each species. Lower πN/πS ratios indicate more efficient purifying selection. First, we generated genome annotations by mapping gene models of zebra finch (RefSeq accession: GCF_048771995.1) onto each bird’s reference genome using Liftoff v1.6.1 (60). We then filtered out gene models containing in-frame STOP codons using the Gffread utility (-V option) from Cufflinks v2.2.1 (61). Using these refined annotations, we identified synonymous and non-synonymous nucleotide substitutions in coding regions for each species with SnpEff v4.3.68 (62). The input biallelic VCF datasets were prepared as follows: raw variants were called from BAM files using Sambamba, and the resulting VCF files were filtered in VCFtools v0.1.17 (63) with the following criteria: -f “Qual=20/MinMQ=30” and -min-alleles 2 -max-alleles 2 -remove-indels - maf 0.1 -max-missing 1.0. Finally, we computed *π*_N_ and *π*_S_ in non-overlapping 10- Mbp windows using vcftools. We note that due to the discrete distribution of coding regions across the genome, the *π*_N_/*π*_S_ ratio was sensitive to window size, decreasing steadily as window size increased up to 10 Mbp (*SI Appendix*, Fig. S6).

### Ecological and life-history variables

To examine the predictive effect of life history on across-species variation of genetic diversity, we compiled four continuous variables indicating reproductive strategy from public databases (58, 64, 65), including maximum longevity, clutch size, adult survival, and body mass.

Environmental factors (e.g., temperature) are known to covary with elevation (66). Here, we used the mean breeding elevations of the HHMs for each species compiled from public sources (*SI Appendix*, Table S1) (67–70) as a proxy for modern environmental conditions.

## Statistical analysis

To account for phylogenetic non-independence, we used MCMCglmm analysis from the R package MCMCglmm (21) to investigate potential drivers of genetic diversity variation in the studied HHM songbirds. Prior to the analysis, we assessed the normality of all explanatory variables using the Shapiro–Wilk test, a robust method for evaluating normal distributions (71). Elevation followed a normal distribution (*p* = 0.38) and was used untransformed, whereas all other variables showing non-normality (*p* < 0.05) were log₁₀-transformed. Tajima’s *D* values were log-transformed after adding a constant of 1.2 to accommodate negative values. All explanatory variables were standardized to a mean of zero and a standard deviation of one to allow direct comparison of effect sizes. These variables were included as fixed effects, with species and phylogeny specified as random effects. The phylogenetic tree was constructed as a consensus from 100 randomly selected trees based on the Ericson backbone (72) using TreeAnnotator in BEAST 2.6.4 (73). We fitted a bivariate-response model with log-transformed *π* and *h* as response variables, and performed 55,000 Markov chain Monte Carlo (MCMC) iterations with a burn-in of 5,000 and a thinning interval of 50. In line with Hadfield (74), we fixed the covariance structure and used noninformative inverse Wishart priors with V = 1 and ν = 0.002 for the variance components. Model convergence was evaluated by inspecting trace plots (*SI Appendix*, Fig. S7) and ensuring low autocorrelation levels (< 0.1, assessed using the ‘autocorr’ function). Collinearity among predictors was assessed via variance inflation factors (VIF), all of which were below the recommended threshold of 10 (75). We further conducted a posterior predictive check using 1,000 simulations under the same model parameters (via the “simulate” function) to confirm that observed means fell within the 95% credible intervals of the posterior predictive distribution for both *π* and *h* (*SI Appendix*, Fig. S8). All analyses were performed in R version 4.3.2 (76).

## Data availability

The scripts used in the paper are deposited in the Science Data Bank (doi: 10.57760/sciencedb.17191). Genomic data have been archived in Genome Sequence Archive (the accession number: CRA020079).

## Acknowledgments

We thank Ri Liu, Ning Cui, Xuerong Li and Xurui Si (Kunming Institute of Zoology, Chinese Academy of Sciences) for assistance in data analysis, and Heqi Wu and Jianyun Gao (Kunming Institute of Zoology, Chinese Academy of Sciences) for sample collection. This work was supported by the National Key R&D Program of China (2022YFC2602500), the National Natural Science Foundation of China (32170440), the Yunnan Applied Basic Research Project (202401AS070078) and the West Light Foundation of the Chinese Academy of Sciences.

## Author contributions

FD conceived the ideas and designed the study; LYD and YRH performed the analyses; FD, FW and YD contributed to collecting the samples; FD led the writing, with substantial input from CMH, which was approved by all the other authors.

## Competing Financial Interests statement

The authors declare no competing financial interests.

## Notes

### Competing Interest Statement

The authors have declared no competing interest.

