## Supplementary material for "Pleistocene demographic histories dominate contemporary genomic diversity in a continental radiation of Himalayan-Hengduan songbirds": Figures S1 to S8 and Tables S1 to S2

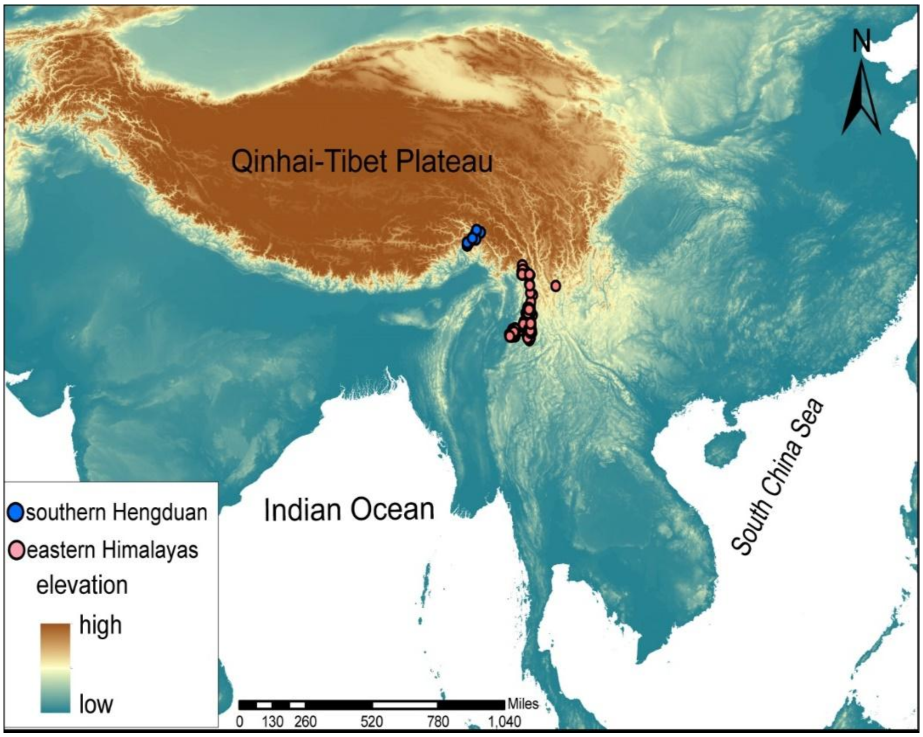


**Fig. S1.** Map of sampling localities in the Himalayan–Hengduan Mountains. The blue circles indicate sampling sites from Nanga Bawa Mountain in the eastern Himalayas, whereas the red circles indicate those from Gaoligong Mountain in the southern Hengduan Mountains. The inset shows the relative location of the study site in southern Asia, with lighter colors indicating higher altitudes.

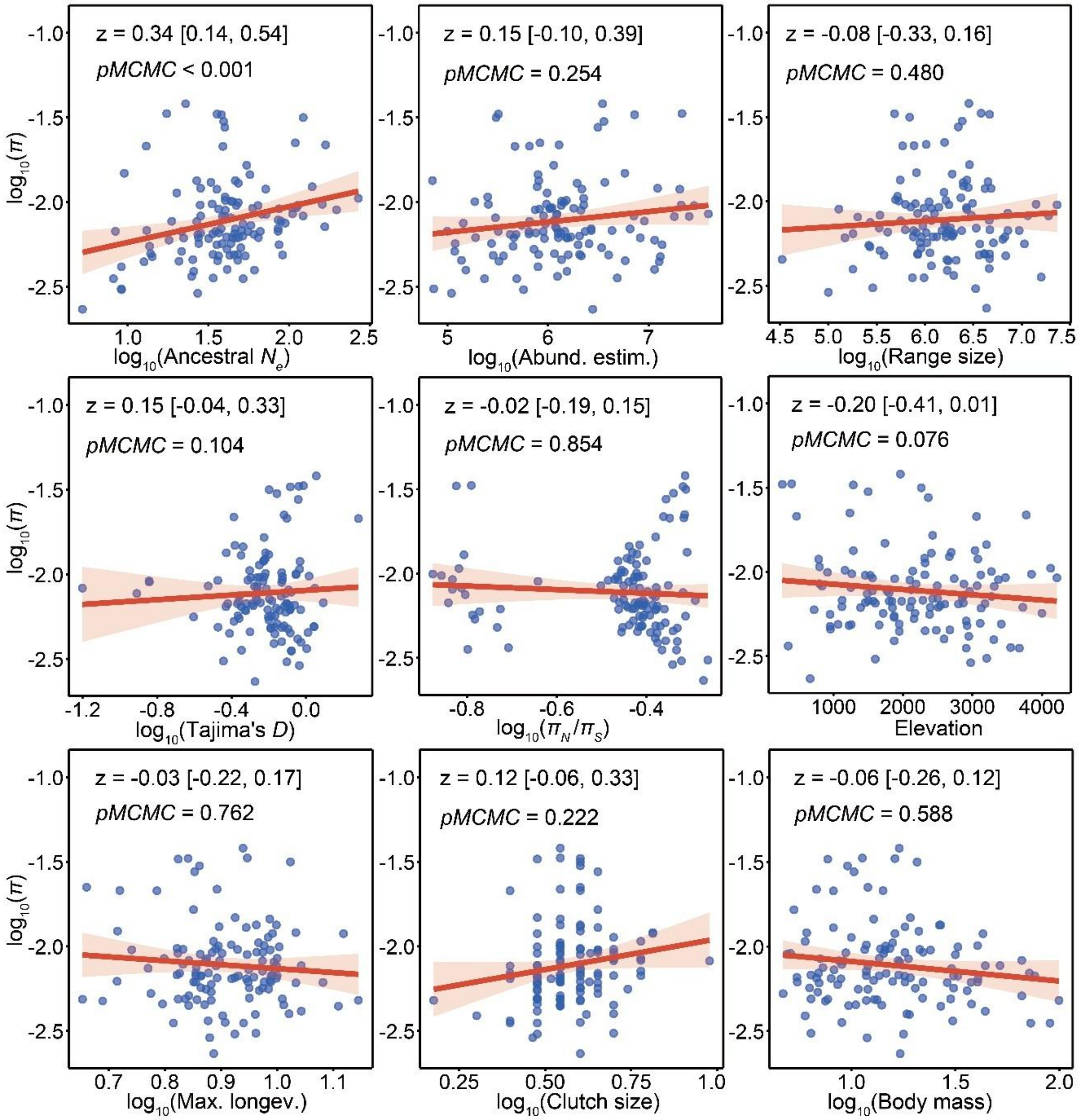


**Fig. S2.** Correlations of genome-wide nucleotide diversity ($\pi$) across the 120 studied birds with potential explanatory variables. These variables include long-term $N_e$, population abundance estimate (Pop. abund.), range size, Tajima's $D$, natural selection (represented by $\pi_N/\pi_S$), elevation, and life-history traits (maximum longevity, Max. longev.; clutch size; and body mass).

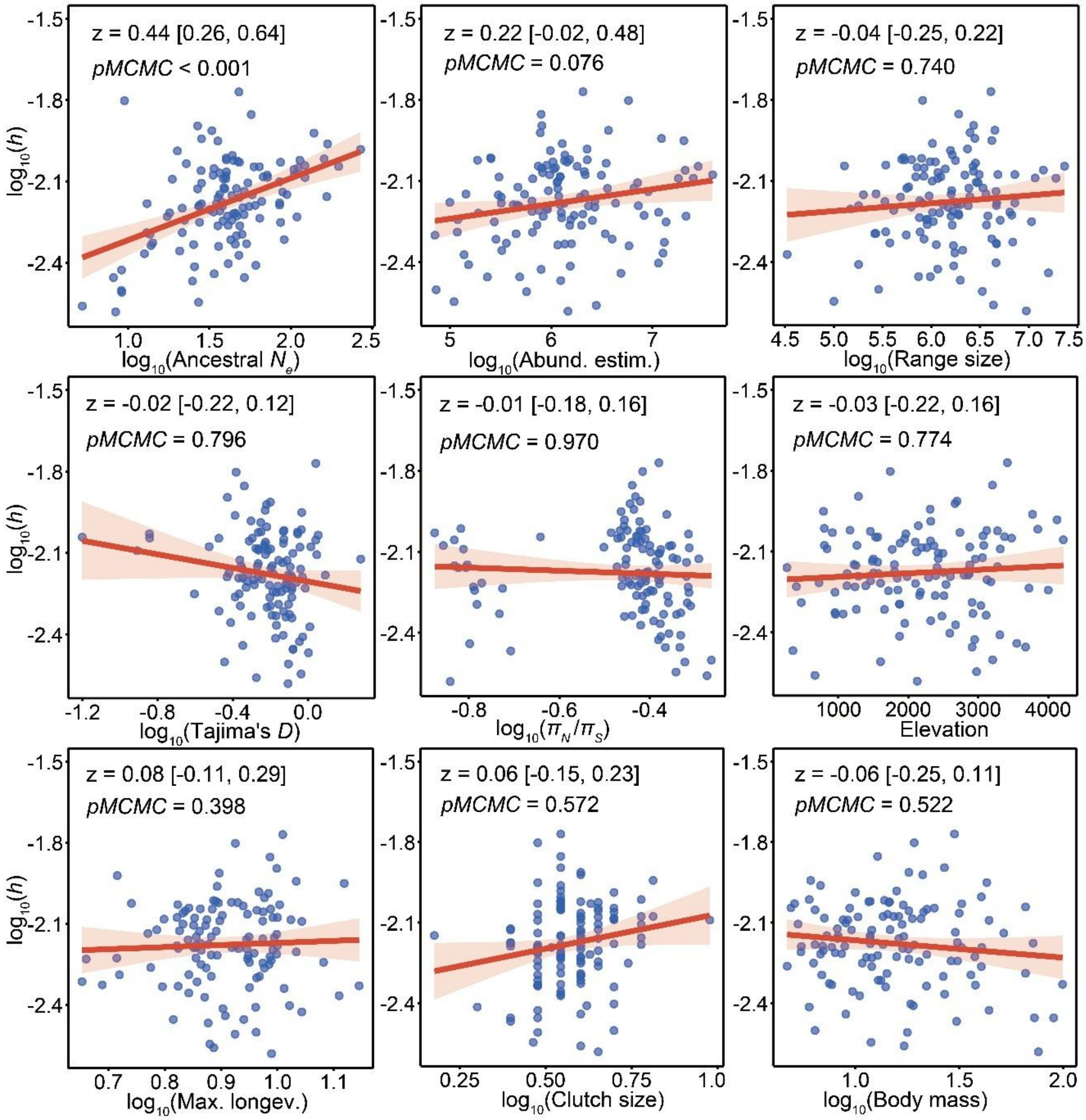


**Fig. S3.** Correlations of genome-wide nucleotide diversity ($\pi$) across the 120 studied birds with potential explanatory variables. These variables include long-term $N_e$, population abundance estimate (Pop. abund.), range size, Tajima's $D$, natural selection (represented by $\pi_N/\pi_S$), elevation, and life-history traits (maximum longevity, Max. longev.; clutch size; and body mass).

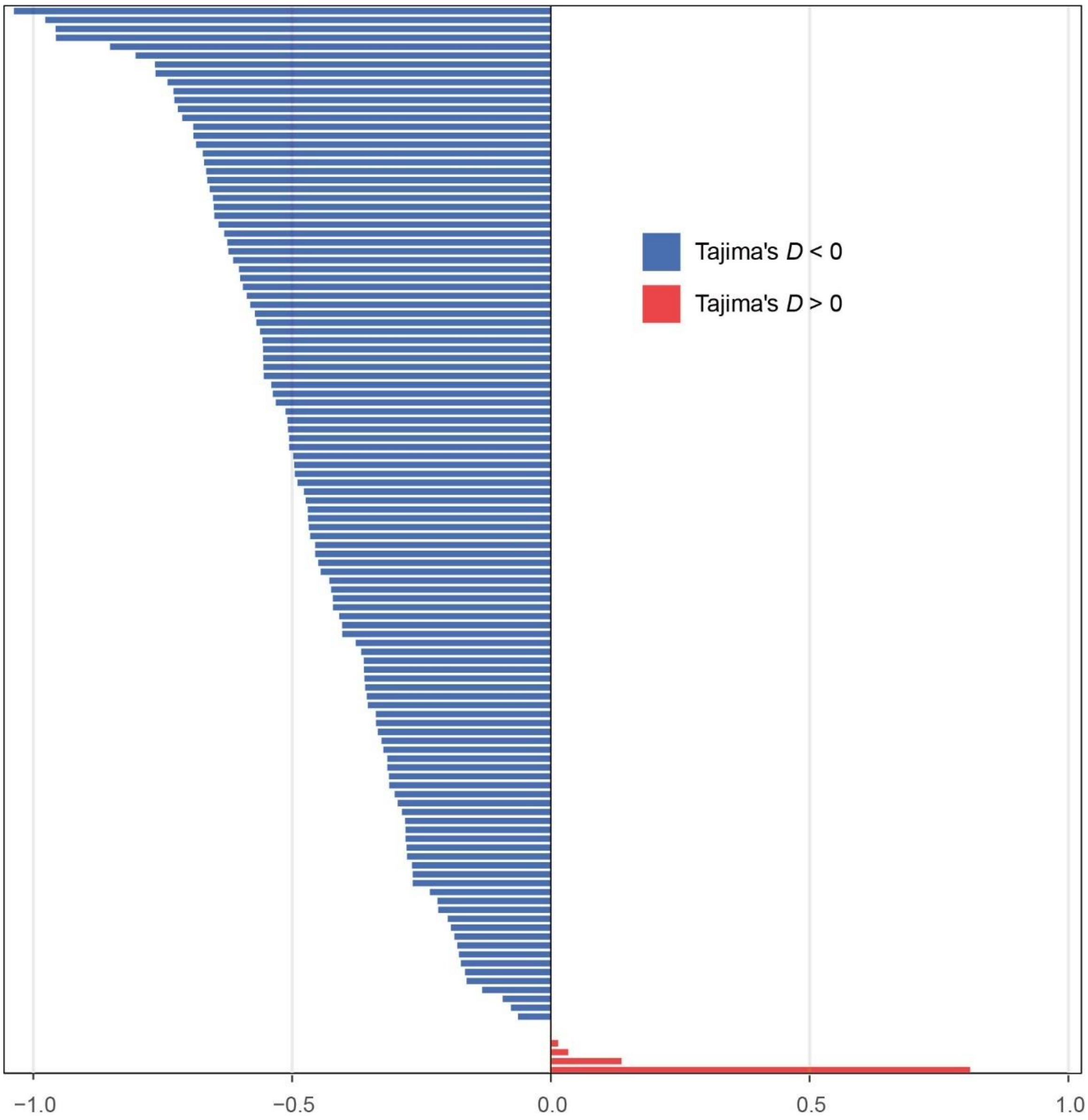


**Fig. S4.** Distribution of Tajima's *D* statistics across the 120 studied songbirds.

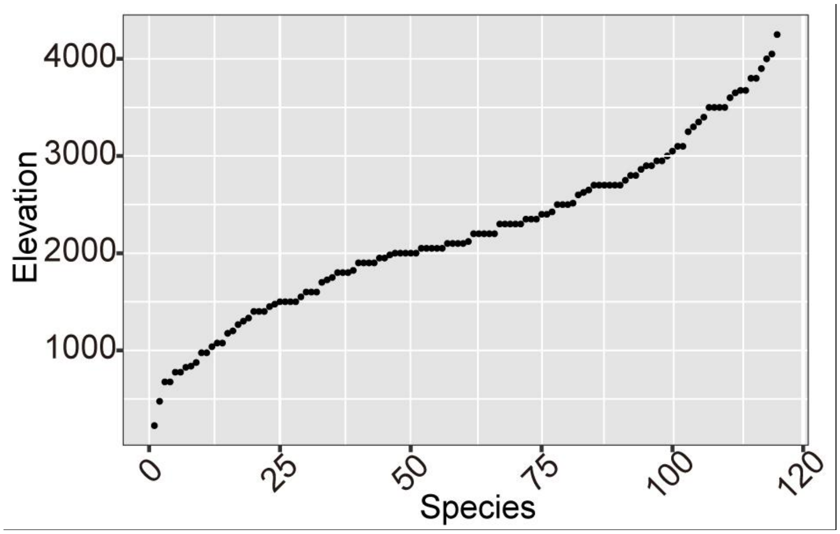


**Fig. S5.** Distribution of midpoint elevation range across the 120 studied songbirds.

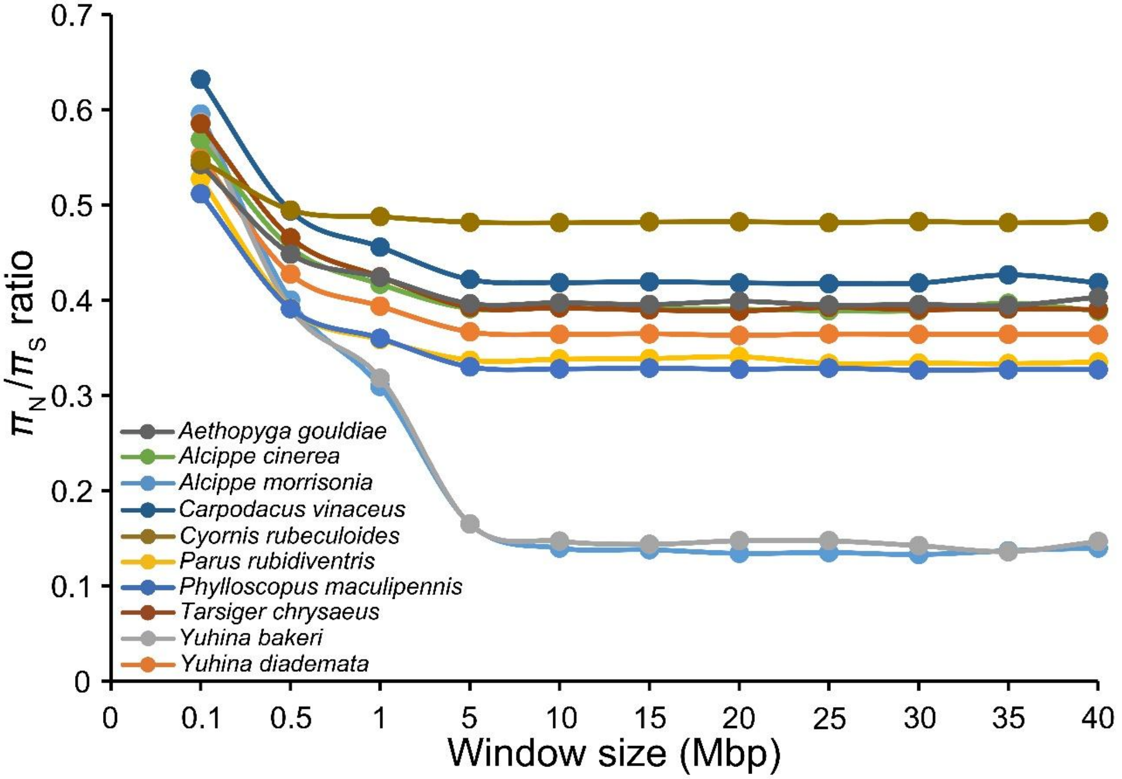


**Fig. S6. Sensitivity of $\pi_N/\pi_S$ ratio to window size.** Analysis based on 10 randomly selected songbird species shows a consistent decrease in $\pi_N/\pi_S$ ratio as window size increases, up to 10 Mbp.

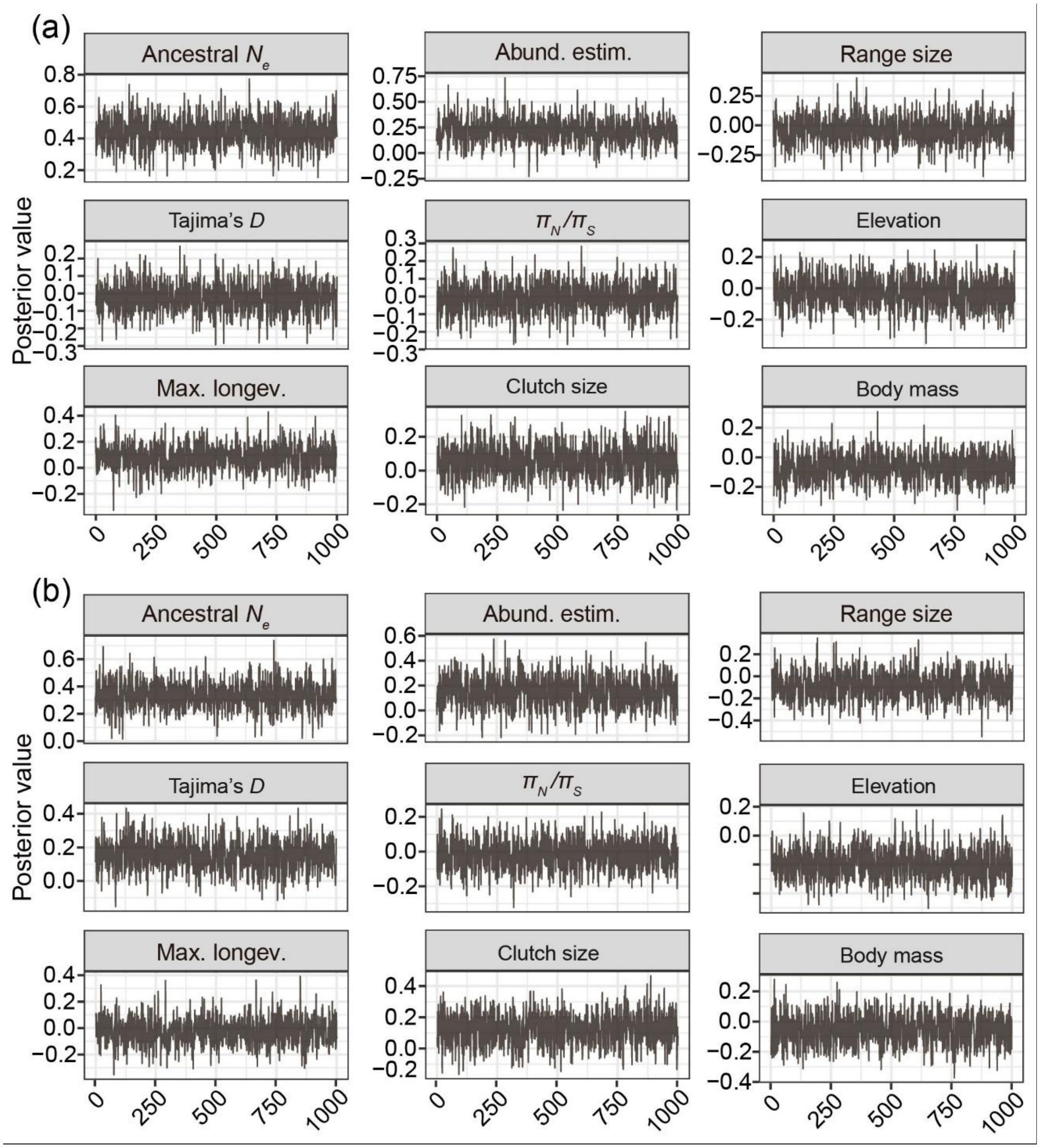


**Fig. S8.** Trace plots of explanatory variables from MCMCglmm simulations for (a) nucleotide diversity and (b) heterogeneity. These variables include long-term $N_e$, population abundance estimate (Pop. abund.), range size, Tajima's *D*, natural selection (represented by $\pi_N/\pi_S$), elevation, and life-history traits (maximum longevity, Max. longev.; clutch size; and body mass).

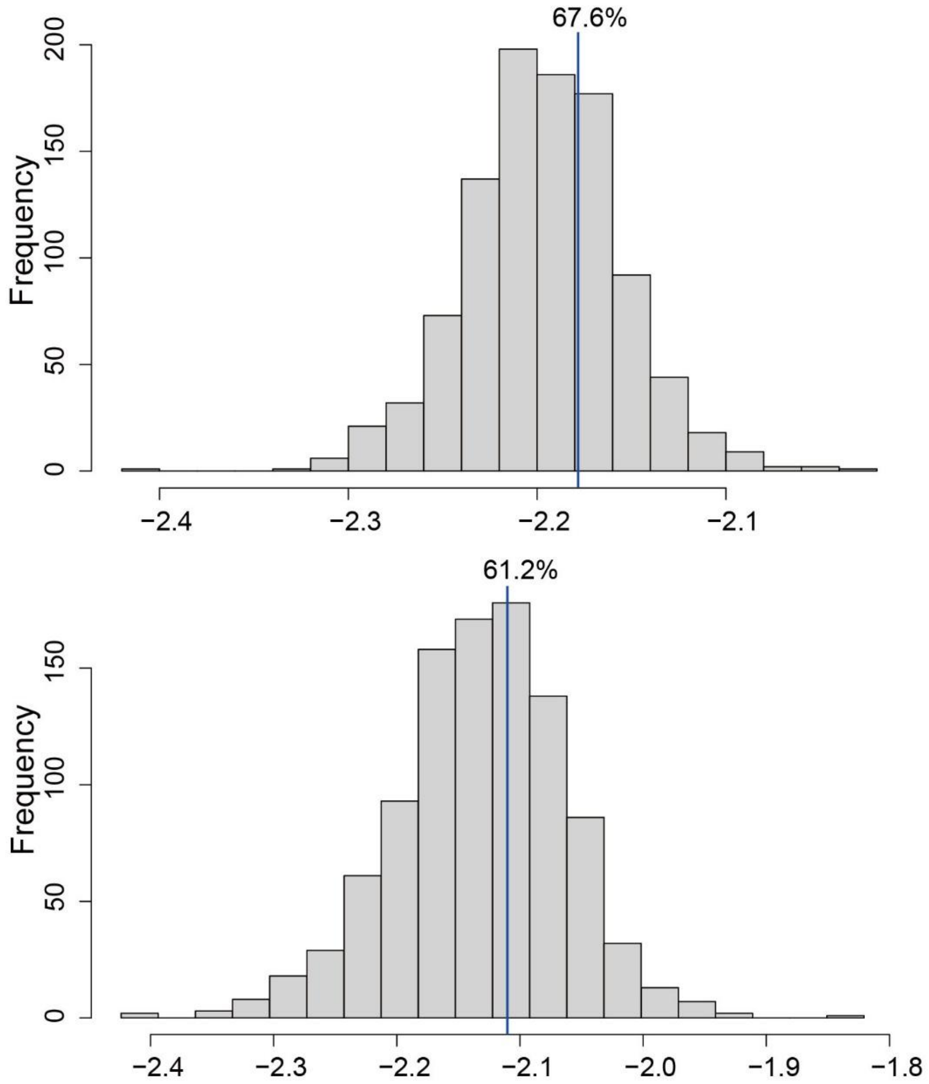


**Fig. S8.** Posterior predictive histograms with 1,000 simulations. The blue line represents the observed value for both nucleotide diversity (a) and heterogeneity (b) and possibility of the observed value ≥ simulated one is indicated above.

**Tables**

**Table S1.** Sampling and sequencing information for reference genomes of the 120 songbirds in the present study.

| Species name | Individual ID | Sampling localities | Sampling time | Sampling latitude | Sampling longitude | sequencing coverage | Scaffold N50 (Mbp) | Contig N50 (Kbp) | Tajima's *D* | BUSCO score | Mean distance (km) | gap | Elevation[citation] |
|---|---|---|---|---|---|---|---|---|---|---|---|---|---|
| *Abroscopus schisticeps* | GLGS1355 | southern Hengduan | 2003 | 95.17747 | 29.24492 | 40.7 | 75.2 | 13.7 | -0.69084 | 94.10% | 196.87 | 7.39% | 2300[1] |
| *Actinodura egertoni* | GLGS5798 | southern Hengduan | 2005 | 95.17747 | 29.24492 | 49.5 | 73.0 | 23.8 | -0.66382 | 95.40% | 241.45 | 6.09% | 1500[1] |
| *Aegithalos concinnus* | GLGS1448 | southern Hengduan | 2003 | 95.17747 | 29.24492 | 40.7 | 91.2 | 6.9 | -0.50764 | 90.00% | 243.74 | 14.39% | 1900[1] |
| *Aegithalos iouschistos* | MT267 | eastern Himalayas | 2018 | 95.17747 | 29.24492 | 49.8 | 80.0 | 8.0 | -0.62515 | 92.00% | 1.25 | 9.48% | 2950[1] |
| *Aethopyga gouldiae* | MT685 | eastern Himalayas | 2020 | 95.17747 | 29.24492 | 41.4 | 95.6 | 7.7 | -0.50910 | 91.40% | 89.17 | 15.99% | 2500[1] |
| *Aethopyga ignicauda* | GLGS6221 | southern Hengduan | 2006 | 95.94088 | 29.7737 | 43.8 | 78.2 | 14.7 | 0.81011 | 95.50% | 1.38 | 8.24% | 3500[1] |
| *Aethopyga nipalensis* | 05216 | southern Hengduan | 2005 | 95.94155 | 29.77433 | 41.8 | 76.7 | 19.7 | -0.40870 | 96.20% | 73.52 | 7.32% | 2400[1] |
| *Alcippe castaneceps* | 04157 | southern Hengduan | 2004 | 95.94088 | 29.7737 | 33.1 | 78.7 | 16.4 | -0.58090 | 95.20% | 89.87 | 8.02% | 2515[2] |
| *Alcippe chrysotis* | GLGS2384 | southern Hengduan | 2004 | 95.9426 | 29.77394 | 65.4 | 78.1 | 15.2 | -0.16582 | 95.30% | 201.45 | 7.18% | 2625[2] |
| *Alcippe cinerea* | 04365 | southern Hengduan | 2004 | 95.94155 | 29.77433 | 35.9 | 82.5 | 13.6 | -0.31589 | 94.70% | 0.00 | 9.97% | 2350[3] |
| *Alcippe dubia* | GLGS1427 | southern Hengduan | 2003 | 95.6484 | 29.72893 | 41.5 | 77.8 | 13.0 | -0.58735 | 93.60% | 281.04 | 8.61% | 1400[3] |
| *Alcippe ludlowi* | MT069 | eastern Himalayas | 2018 | 95.70076 | 29.8753 | 48 | 76.0 | 20.3 | -0.07729 | 96.10% | 6.91 | 5.70% | 3500[4] |

| | | | | | | | | | | | | | |
|---|---|---|---|---|---|---|---|---|---|---|---|---|---|
| *Alcippe morrisonia* | MT156 | eastern Himalayas | 2018 | 95.70112 | 29.87534 | 44.2 | 83.6 | 12.8 | -0.06341 | 93.90% | 39.81 | 9.29% | 1265[2] |
| *Alcippe vinipectus* | GLGS2238 | southern Hengduan | 2004 | 95.6484 | 29.72893 | 33.3 | 79.0 | 14.6 | -0.32728 | 95.10% | 1.93 | 8.43% | 2862.5[2] |
| *Anthus hodgsoni* | 2022LS102 | southern Hengduan | 2022 | 95.47852 | 29.49969 | 42.2 | 89.0 | 7.9 | -0.50576 | 90.70% | 198.17 | 13.90% | 2800[1] |
| *Arachnothera magna* | MT666 | eastern Himalayas | 2020 | 95.17699 | 29.24506 | 52.5 | 79.4 | 21.9 | -0.09287 | 95.90% | 0.41 | 6.61% | 1075[1] |
| *Carpodacus edwardsii* | MT248 | eastern Himalayas | 2018 | 95.17765 | 29.24599 | 48.1 | 77.2 | 18.0 | -0.49482 | 96.30% | 123.30 | 7.01% | 3650[1] |
| *Carpodacus erythrinus* | MT221 | eastern Himalayas | 2018 | 95.17816 | 29.24692 | 44.9 | 87.0 | 10.6 | -0.59502 | 93.40% | 77.19 | 11.78% | 3675[1] |
| *Carpodacus nipalensis* | MT411 | eastern Himalayas | 2020 | 95.17741 | 29.12477 | 36.4 | 89.3 | 10.1 | -0.35564 | 93.00% | 108.40 | 14.06% | 3900[1] |
| *Carpodacus thura* | MT005 | eastern Himalayas | 2018 | 95.4551 | 29.47404 | 46 | 81.7 | 10.8 | -0.44941 | 94.30% | 26.05 | 10.50% | 3800[1] |
| *Carpodacus vinaceus* | GLG23158 | southern Hengduan | 2023 | 95.69757 | 29.80727 | 48.6 | 84.0 | 13.1 | -0.47722 | 94.60% | 271.18 | 10.63% | 3100[1] |
| *Certhia familiaris* | MT020 | eastern Himalayas | 2018 | 95.69757 | 29.80727 | 49 | 76.3 | 13.2 | -0.65022 | 94.90% | 37.00 | 6.50% | 3350[1] |
| *Cettia flavolivacea* | GLGS5279 | southern Hengduan | 2005 | 95.69757 | 29.80727 | 44.4 | 87.2 | 5.7 | -0.33440 | 90.20% | 237.92 | 12.85% | 2600[1] |
| *Cettia fortipes* | MT656 | eastern Himalayas | 2020 | 95.94032 | 29.77333 | 53.4 | 76.6 | 20.8 | -0.40338 | 96.50% | 117.98 | 5.77% | 2100[1] |
| *Chaimarrornis leucocephalus* | MT065 | eastern Himalayas | 2018 | 95.69757 | 29.80727 | 52.3 | 76.5 | 9.9 | 0.00141 | 93.30% | 52.51 | 8.29% | 2950[1] |
| *Chelidorhynx hypoxantha* | GLGS1420 | southern Hengduan | 2003 | 95.17722 | 29.24925 | 50.5 | 74.2 | 12.8 | -0.42779 | 94.10% | 270.91 | 6.84% | 2700[1] |
| *Chrysomma sinense* | GLG23673 | southern Hengduan | 2023 | 95.17894 | 29.24925 | 49.7 | 73.9 | 16.2 | -0.56895 | 96.10% | 239.97 | 5.57% | 225[1] |
| *Cinclidium leucurum* | GLGS5775 | southern Hengduan | 2005 | 95.24837 | 29.24837 | 43.7 | 84.2 | 12.8 | -0.72758 | 95.10% | 189.02 | 8.40% | 1822.5[2] |
| *Cinclus pallasii* | GLGS5810 | southern Hengduan | 2005 | 95.17588 | 29.24723 | 40.8 | 72.3 | 20.7 | -0.31589 | 97.20% | 241.68 | 4.43% | 2300[1] |

| | | | | | | | | | | | | | |
|---|---|---|---|---|---|---|---|---|---|---|---|---|---|
| *Culicicapa ceylonensis* | GLGS6286 | southern Hengduan | 2006 | 95.17455 | 29.24654 | 42.9 | 82.1 | 9.5 | -0.27799 | 93.20% | 248.45 | 9.82% | 1725[1] |
| *Cyornis banyumas* | MT105 | eastern Himalayas | 2018 | 95.69849 | 29.806 | 67.2 | 76.5 | 14.6 | -0.55631 | 95.10% | 30.72 | 7.01% | 1600[3] |
| *Cyornis poliogenys* | MT448 | eastern Himalayas | 2020 | 95.69849 | 29.806 | 46.6 | 80.0 | 10.1 | -0.18647 | 92.50% | 32.19 | 10.58% | 775[1] |
| *Cyornis rubeculoides* | NJDL001 | southern Hengduan | 2017 | 95.64767 | 29.72748 | 48.7 | 76.3 | 11.8 | -0.31243 | 94.30% | 264.66 | 7.67% | 675[1] |
| *Dicrurus leucophaeus* | 2022GS281 | southern Hengduan | 2022 | 95.6484 | 29.72893 | 50.1 | 79.2 | 20.0 | -0.63099 | 96.10% | 268.56 | 6.97% | 1900[1] |
| *Emberiza elegans* | GLGS1464 | southern Hengduan | 2003 | 95.69859 | 29.80631 | 44.4 | 84.1 | 11.3 | 0.01413 | 92.60% | 274.43 | 11.72% | 2300[3] |
| *Enicurus maculatus* | GLGS5185 | southern Hengduan | 2005 | 95.17319 | 29.24741 | 50.5 | 77.7 | 9.9 | -0.28085 | 93.40% | 288.96 | 8.69% | 1700[1] |
| *Enicurus scouleri* | GLGS2400 | southern Hengduan | 2004 | 95.17319 | 29.24741 | 41.5 | 95.8 | 9.1 | -0.31296 | 89.70% | 140.12 | 8.39% | 2100[1] |
| *Erpornis zantholeuca* | DHLC004 | southern Hengduan | 2014 | 95.17464 | 29.24826 | 38.8 | 80.5 | 13.2 | -0.45535 | 94.00% | 223.09 | 8.46% | 825[1] |
| *Eumyias thalassinus* | GLGS6424 | southern Hengduan | 2006 | 95.17464 | 29.24826 | 42.4 | 89.9 | 6.5 | -0.46958 | 90.10% | 282.63 | 13.79% | 1550[1] |
| *Ficedula albicilla* | MT261 | eastern Himalayas | 2018 | 95.17455 | 29.24654 | 47 | 81.1 | 9.9 | -0.85143 | 92.90% | 121.14 | 11.29% | 1332.5[3] |
| *Ficedula hodgsonii* | GLGS1803 | southern Hengduan | 2003 | 95.17672 | 29.24811 | 51.3 | 95.8 | 5.6 | -0.53989 | 88.70% | 227.80 | 15.62% | 1037.5[2] |
| *Ficedula hyperythra* | MT680 | eastern Himalayas | 2020 | 95.17811 | 29.24603 | 50.4 | 83.7 | 10.4 | -0.65279 | 92.90% | 3.10 | 9.93% | 2300[1] |
| *Ficedula strophiata* | MT058 | eastern Himalayas | 2018 | 95.17586 | 29.24616 | 44.2 | 85.9 | 8.4 | -0.64189 | 90.90% | 5.94 | 12.37% | 3050[1] |
| *Ficedula tricolor* | GLGS5173 | southern Hengduan | 2005 | 95.17641 | 29.24724 | 39.3 | 79.3 | 9.9 | -0.56183 | 92.70% | 202.14 | 10.66% | 2425[1] |
| *Garrulax affinis* | MT034 | eastern Himalayas | 2018 | 95.17565 | 29.24814 | 48.8 | 77.4 | 19.8 | -0.26680 | 95.90% | 129.31 | 6.02% | 2050[2] |
| *Garrulax erythrocephalus* | MT275 | eastern Himalayas | 2018 | 95.4829 | 29.50108 | 44 | 75.5 | 21.8 | -0.44474 | 95.00% | 3.48 | 6.33% | 2050[2] |

| | | | | | | | | | | | | | |
|---|---|---|---|---|---|---|---|---|---|---|---|---|---|
| *Garrulax subunicolor* | GLGS2167 | southern Hengduan | 2004 | 95.47757 | 29.49919 | 44.9 | 76.9 | 12.9 | -0.40294 | 93.30% | 233.71 | 8.03% | 3675[2] |
| *Hemixos flavala* | MT541 | eastern Himalayas | 2020 | 95.47757 | 29.49919 | 49.3 | 81.8 | 14.8 | -0.21743 | 95.30% | 16.30 | 8.70% | 775[1] |
| *Heterophasia desgodinsi* | GLGS6416 | southern Hengduan | 2006 | 95.45539 | 29.47474 | 45 | 77.9 | 14.4 | -0.72069 | 93.70% | 243.46 | 8.71% | 2050[3] |
| *Heterophasia pulchella* | 04036 | southern Hengduan | 2004 | 95.17665 | 29.2464 | 36.8 | 77.1 | 13.8 | -0.65144 | 93.10% | 209.97 | 8.30% | 2400[1] |
| *Hypsipetes leucocephalus* | MT671 | eastern Himalayas | 2020 | 95.18019 | 29.2486 | 46.3 | 83.7 | 12.0 | -0.55709 | 94.40% | 0.31 | 12.07% | 1475[1] |
| *Hypsipetes mcclellandii* | MT582 | eastern Himalayas | 2020 | 95.18019 | 29.2486 | 49.5 | 84.3 | 12.9 | -0.67258 | 95.00% | 60.65 | 10.09% | 1600[2] |
| *Leiothrix argentauris* | MT663 | eastern Himalayas | 2020 | 95.17665 | 29.2464 | 52.3 | 77.4 | 22.7 | -0.42447 | 95.70% | 6.92 | 6.06% | 1200[1] |
| *Leiothrix lutea* | GLGS5253 | southern Hengduan | 2005 | 95.17665 | 29.2464 | 35.8 | 106.4 | 3.9 | -0.68539 | 86.70% | 21.54 | 16.64% | 2500[1] |
| *Lonchura punctulata* | MT134 | eastern Himalayas | 2018 | 95.17665 | 29.2464 | 39.2 | 98.2 | 6.3 | -0.80235 | 88.60% | 31.69 | 17.84% | 2350[1] |
| *Minla cyanouroptera* | GLGS6485 | southern Hengduan | 2006 | 95.1775 | 29.24667 | 48.3 | 74.2 | 11.4 | 0.03357 | 92.80% | 247.24 | 7.39% | 2000[1] |
| *Minla ignotincta* | 04469 | southern Hengduan | 2004 | 95.17808 | 29.24648 | 34.6 | 79.9 | 10.0 | -0.51287 | 91.00% | 243.38 | 10.64% | 2500[1] |
| *Minla strigula* | GLGS5100 | southern Hengduan | 2005 | 95.17539 | 29.24321 | 49.7 | 79.7 | 13.5 | -0.28783 | 94.00% | 202.26 | 8.52% | 2900[1] |
| *Myzornis pyrrhoura* | MT107 | eastern Himalayas | 2018 | 95.17905 | 29.24533 | 50.6 | 76.1 | 19.6 | -0.18057 | 96.50% | 52.94 | 5.11% | 3300[1] |
| *Niltava grandis* | GLGS1463 | southern Hengduan | 2003 | 95.17893 | 29.14447 | 46.5 | 75.8 | 6.5 | -0.36646 | 83.70% | 140.94 | 13.66% | 1900[1] |
| *Niltava macgrigoriae* | MT132 | eastern Himalayas | 2018 | 95.45462 | 29.47416 | 48.6 | 76.2 | 15.8 | -0.33830 | 95.90% | 79.88 | 6.61% | 1400[1] |
| *Niltava sundara* | MT136 | eastern Himalayas | 2018 | 95.45462 | 29.47416 | 47 | 81.8 | 6.3 | -0.49773 | 88.90% | 98.85 | 13.43% | 2300[1] |
| *Orthotomus sutorius* | MT526 | eastern Himalayas | 2020 | 95.44713 | 29.48116 | 52.4 | 78.7 | 24.4 | -0.30181 | 97.10% | 3.88 | 4.75% | 975[1] |

| | | | | | | | | | | | | | |
|---|---|---|---|---|---|---|---|---|---|---|---|---|---|
| *Paradoxornis fulvifrons* | GLGS0352 | southern Hengduan | 2002 | 95.44713 | 29.48116 | 49 | 76.3 | 12.7 | -0.74070 | 94.80% | 203.63 | 6.56% | 3100[1] |
| *Paradoxornis nipalensis* | GLG04531 | southern Hengduan | 2005 | 95.64801 | 29.72855 | 45.2 | 77.4 | 10.4 | -0.23369 | 94.60% | 52.54 | 9.58% | 2200[1] |
| *Parus ater* | GLGS0434 | eastern Himalayas | 2018 | 95.6484 | 29.72893 | 55.2 | 83.1 | 9.3 | -0.48944 | 94.30% | 18.26 | 11.65% | 3400[1] |
| *Parus monticolus* | MT137 | eastern Himalayas | 2018 | 95.6484 | 29.72893 | 49.3 | 75.8 | 21.6 | -0.27883 | 95.90% | 69.58 | 6.07% | 1950[1] |
| *Parus rubidiventris* | MT004 | eastern Himalayas | 2018 | 95.64763 | 29.72734 | 53.3 | 77.4 | 18.8 | -0.46514 | 96.50% | 83.59 | 6.82% | 3500[1] |
| *Parus spilonotus* | MT536 | eastern Himalayas | 2020 | 95.64801 | 29.72855 | 48.3 | 81.5 | 17.1 | -0.55475 | 95.50% | 171.26 | 9.40% | 1800[1] |
| *Passer rutilans* | MT170 | eastern Himalayas | 2018 | 95.6484 | 29.72893 | 46.3 | 76.1 | 18.6 | -0.36135 | 95.90% | 13.64 | 6.85% | 2200[1] |
| *Phoenicurus frontalis* | MT003 | eastern Himalayas | 2018 | 95.64802 | 29.72855 | 45.4 | 82.0 | 5.8 | -0.60252 | 89.10% | 67.02 | 14.39% | 3800[1] |
| *Phoenicurus schisticeps* | MT313 | eastern Himalayas | 2018 | 95.64802 | 29.72855 | 51.4 | 77.9 | 15.5 | -0.28085 | 94.20% | 11.59 | 8.31% | 4250[1] |
| *Phylloscopus affinis* | MT642 | eastern Himalayas | 2020 | 95.47757 | 29.49919 | 46.7 | 79.2 | 9.8 | -0.55593 | 93.20% | 1.64 | 9.37% | 4050[1] |
| *Phylloscopus armandii* | 2022GS206 | southern Hengduan | 2022 | 95.47757 | 29.49919 | 44.4 | 86.9 | 7.2 | -0.76480 | 91.70% | 299.98 | 13.21% | 1450[3] |
| *Phylloscopus cantator* | MT530 | eastern Himalayas | 2020 | 95.6484 | 29.72893 | 46.9 | 87.5 | 12.9 | -0.17767 | 93.20% | 28.40 | 9.53% | 875[1] |
| *Phylloscopus chloronotus* | MT002 | eastern Himalayas | 2018 | 95.454 | 29.47356 | 38.9 | 95.6 | 2.1 | -0.95635 | 77.40% | 19.58 | 22.04% | 3000[1] |
| *Phylloscopus davisoni* | 2022GS048 | southern Hengduan | 2022 | 95.454 | 29.47356 | 41.2 | 88.7 | 5.4 | 0.00142 | 91.20% | 278.60 | 12.88% | 2050[3] |
| *Phylloscopus fuscatus* | MT138 | eastern Himalayas | 2018 | 95.454 | 29.47356 | 49.9 | 76.5 | 15.2 | -0.55545 | 95.80% | 42.30 | 6.62% | 837.5[2] |
| *Phylloscopus maculipennis* | 04190 | southern Hengduan | 2004 | 95.454 | 29.47356 | 39 | 81.2 | 8.2 | -0.95676 | 91.50% | 205.31 | 10.86% | 2700[1] |
| *Phylloscopus magnirostris* | MT625 | eastern Himalayas | 2020 | 95.454 | 29.47356 | 47.6 | 82.9 | 9.6 | -0.53727 | 92.80% | 52.10 | 11.24% | 2700[1] |

| | | | | | | | | | | | | | |
|---|---|---|---|---|---|---|---|---|---|---|---|---|---|
| *Phylloscopus pulcher* | MT094 | eastern Himalayas | 2018 | 95.17698 | 29.24691 | 42.7 | 87.8 | 8.1 | -0.69100 | 92.10% | 80.07 | 13.22% | 2000[1] |
| *Phylloscopus xanthoschistos* | MT133 | eastern Himalayas | 2018 | 95.18027 | 29.24741 | 46.7 | 79.5 | 13.0 | -0.46985 | 94.80% | 24.48 | 8.36% | 1500[1] |
| *Pnoepyga pusilla* | MT498 | eastern Himalayas | 2020 | 95.18277 | 29.24446 | 51.9 | 77.7 | 18.9 | -0.47352 | 96.70% | 22.56 | 6.84% | 1800[1] |
| *Pomatorhinus ruficollis* | GLG23135 | southern Hengduan | 2023 | 95.18277 | 29.24446 | 72.4 | 74.3 | 16.5 | -0.36041 | 95.40% | 247.70 | 8.04% | 2000[2] |
| *Prinia atrogularis* | MT643 | eastern Himalayas | 2020 | 95.1757 | 29.25349 | 51.4 | 70.9 | 26.7 | -0.62309 | 97.40% | 180.33 | 3.49% | 1500[1] |
| *Prinia hodgsonii* | GLG23691 | southern Hengduan | 2023 | 95.94086 | 29.77398 | 47.9 | 81.4 | 14.7 | 0.13656 | 94.50% | 251.70 | 8.36% | 475[1] |
| *Prunella immaculata* | MT204 | eastern Himalayas | 2018 | 95.94189 | 29.77446 | 45.1 | 81.3 | 12.3 | -0.46760 | 94.20% | 3.96 | 10.75% | 2650[1] |
| *Prunella strophiata* | MT177 | eastern Himalayas | 2018 | 95.94032 | 29.77333 | 43.4 | 87.0 | 9.0 | -0.61395 | 93.00% | 10.37 | 13.07% | 4000[1] |
| *Pteruthius melanotis* | 04150 | southern Hengduan | 2004 | 95.69757 | 29.80727 | 49.5 | 76.6 | 18.7 | -0.35873 | 96.20% | 320.46 | 5.57% | 1950[1] |
| *Pycnonotus cafer* | MT457 | eastern Himalayas | 2020 | 95.33519 | 29.31587 | 47.4 | 82.3 | 17.2 | -0.17371 | 95.20% | 1.41 | 7.83% | 1075[1] |
| *Pycnonotus flavescens* | GLG23239 | southern Hengduan | 2023 | 95.33519 | 29.31587 | 51.2 | 76.8 | 19.8 | -0.37693 | 96.00% | 230.76 | 6.88% | 1500[3] |
| *Pycnonotus xanthorrhous* | YL20080017 | southern Hengduan | 2008 | 95.17672 | 29.24811 | 49.6 | 75.1 | 15.9 | -0.42102 | 95.40% | 309.49 | 6.42% | 1600[3] |
| *Pyrrhoplectes epauletta* | MT290 | eastern Himalayas | 2018 | 95.17672 | 29.24811 | 47.8 | 79.4 | 16.3 | -0.16253 | 96.00% | 91.37 | 8.24% | 2750[1] |
| *Rhipidura albicollis* | MT594 | eastern Himalayas | 2020 | 95.17672 | 29.24811 | 53.6 | 77.8 | 23.7 | -0.76389 | 96.80% | 112.59 | 5.24% | 1800[1] |
| *Rhyacornis fuliginosa* | 04423 | southern Hengduan | 2004 | 95.66562 | 29.43575 | 38.6 | 75.3 | 9.7 | -0.42130 | 91.40% | 247.95 | 9.07% | 2100[1] |
| *Saxicola ferreus* | MT198 | eastern Himalayas | 2018 | 95.70018 | 29.87567 | 50.3 | 75.2 | 15.5 | -0.66605 | 94.80% | 15.96 | 6.94% | 2200[1] |
| *Seicercus burkii* | MT062 | southern Hengduan | 2018 | 95.70075 | 29.87569 | 40.8 | 78.4 | 12.4 | -0.65910 | 93.60% | 20.62 | 8.94% | 2000[1] |

| | | | | | | | | | | | | | |
|---|---|---|---|---|---|---|---|---|---|---|---|---|---|
| *Seicercus castaniceps* | GLGS2155 | southern Hengduan | 2004 | 95.69992 | 29.87557 | 45.4 | 82.6 | 6.7 | -0.50596 | 90.60% | 214.47 | 11.01% | 2100[1] |
| *Seicercus poliogenys* | 04285 | southern Hengduan | 2004 | 95.70023 | 29.87602 | 46.6 | 78.4 | 12.7 | -0.66991 | 95.10% | 204.19 | 8.68% | 1750[1] |
| *Spizixos canifrons* | GLGS6462 | southern Hengduan | 2006 | 95.45462 | 29.47416 | 36.5 | 99.0 | 10.7 | -0.97695 | 92.80% | 282.56 | 22.37% | 2050[3] |
| *Stachyris chrysaea* | MT641 | eastern Himalayas | 2020 | 97.69839 | 24.63172 | 44.3 | 87.8 | 8.2 | -0.26819 | 91.50% | 6.90 | 13.68% | 2120[2] |
| *Stachyris nigriceps* | MT149 | eastern Himalayas | 2018 | 97.69839 | 24.63172 | 51.4 | 77.4 | 13.6 | -0.28136 | 94.40% | 139.37 | 7.29% | 975[1] |
| *Stachyris ruficeps* | MT148 | eastern Himalayas | 2018 | 97.69839 | 24.63172 | 50.1 | 77.4 | 12.5 | -0.19333 | 93.70% | 14.63 | 8.00% | 1982.5[2] |
| *Sylviparus modestus* | GLGS2190 | southern Hengduan | 2004 | 98.71108 | 25.97792 | 43.8 | 88.9 | 7.5 | -0.57184 | 92.10% | 0.91 | 13.43% | 2700[1] |
| *Tarsiger chrysaeus* | MT035 | eastern Himalayas | 2018 | 98.68167 | 25.97294 | 44.4 | 83.9 | 6.8 | -0.35380 | 90.10% | 77.69 | 14.21% | 3600[1] |
| *Tarsiger indicus* | GLGS5011 | southern Hengduan | 2004 | 97.83313 | 24.95795 | 44 | 87.7 | 6.6 | -0.21919 | 89.70% | 63.41 | 14.19% | 3250[1] |
| *Tephrodornis gularis* | DHRL011 | southern Hengduan | 2014 | 97.75326 | 24.99136 | 57 | 80.2 | 14.9 | -0.60049 | 95.90% | 18.72 | 7.80% | 675[1] |
| *Tesia castaneocoronata* | MT154 | eastern Himalayas | 2018 | 98.62719 | 25.78592 | 47.9 | 78.4 | 14.4 | -0.72909 | 94.60% | 7.70 | 8.10% | 2800[1] |
| *Tesia cyaniventer* | 04130 | southern Hengduan | 2004 | 98.76033 | 24.85558 | 37.8 | 75.1 | 12.0 | -0.71229 | 92.30% | 57.62 | 8.04% | 2000[1] |
| *Turdus albocinctus* | MT072 | eastern Himalayas | 2018 | 98.31179 | 27.68896 | 47.6 | 77.9 | 11.0 | -0.45535 | 93.80% | 155.09 | 8.49% | 2700[1] |
| *Turdus dissimilis* | DHYJ058 | southern Hengduan | 2014 | 98.70558 | 25.98533 | 40.3 | 87.1 | 7.3 | -0.49578 | 90.10% | 42.16 | 13.01% | 1300[3] |
| *Yuhina bakeri* | MT635 | eastern Himalayas | 2020 | 98.71172 | 25.97597 | 48.2 | 78.0 | 17.5 | -0.29584 | 94.70% | 203.34 | 7.82% | 1400[1] |
| *Yuhina castaniceps* | DHYJ129 | southern Hengduan | 2014 | 98.32495 | 27.82627 | 42.7 | 82.3 | 11.2 | -0.36110 | 93.40% | 141.07 | 10.50% | 2350[3] |
| *Yuhina diademata* | GLGS2175 | southern Hengduan | 2004 | 98.65953 | 25.00933 | 42.7 | 75.4 | 18.2 | -0.33785 | 95.70% | 43.37 | 6.33% | 2200[3] |

| | | | | | | | | | | | | | |
|---|---|---|---|---|---|---|---|---|---|---|---|---|---|
| *Yuhina flavicollis* | MT126 | eastern Himalayas | 2018 | 98.71136 | 25.97803 | 50 | 77.9 | 11.2 | -0.53159 | 94.20% | 117.71 | 7.91% | 2200[1] |
| *Yuhina gularis* | MT207 | eastern Himalayas | 2018 | 98.75967 | 24.85608 | 48.2 | 76.6 | 15.3 | -1.03720 | 95.80% | 71.61 | 6.94% | 2700[1] |
| *Yuhina occipitalis* | GLGS1318 | southern Hengduan | 2003 | 98.65925 | 26.00803 | 39.6 | 78.9 | 13.1 | -0.19896 | 95.60% | 0.00 | 7.82% | 2900[1] |
| *Zoothera dixoni* | GLGS1341 | southern Hengduan | 2003 | 98.67897 | 25.97989 | 69.1 | 75.3 | 22.0 | -0.26680 | 96.10% | 111.25 | 4.85% | 3500[1] |
| *Zosterops_japonicus* | MT672 | eastern Himalayas | 2020 | 98.70486 | 25.98536 | 49.5 | 77.7 | 17.0 | -0.13268 | 95.60% | 201.06 | 6.70% | 1900[1] |
| *Zosterops palpebrosus* | MT658 | eastern Himalayas | 2020 | 98.70486 | 25.98536 | 49.9 | 77.6 | 17.8 | -0.32357 | 95.90% | 1.45 | 6.28% | 1175[3] |

**Table S2.** Sampling and sequencing information for population genomes of the 120 songbirds in the present study. Asterisks (*) indicate data originally generated for reference genome assemblies that we downsampled to 30% to reduce the potential confounding effects of an exceedingly high sequencing coverage level on subsequent analyses.

| Species | Individual ID | Sampling localities | Sampling time | Sampling type | Sampling longitude | Sampling latitude | Sequencing coverage |
|---|---|---|---|---|---|---|---|
| *Abroscopus schisticeps* | 2022LS037 | southern Hengduan | 2022 | muscle | 98.61824 | 26.0414 | 15.9 |
| | 2022LS040 | southern Hengduan | 2022 | muscle | 98.61824 | 26.0414 | 16.3 |
| | 2022LS043 | southern Hengduan | 2022 | muscle | 98.61824 | 26.0414 | 17 |
| | 2022LS044 | southern Hengduan | 2022 | muscle | 98.61824 | 26.0414 | 14.1 |
| | GLGS1355* | southern Hengduan | 2003 | muscle | 95.17747 | 29.24492 | 12.2 |
| *Actinodura egertoni* | 2022GS199 | southern Hengduan | 2022 | muscle | 98.79335 | 25.21983 | 13.2 |
| | GLG23286 | southern Hengduan | 2023 | blood | 98.79335 | 25.21985 | 15.5 |
| | GLG23290 | southern Hengduan | 2023 | blood | 98.79335 | 25.21985 | 12.8 |
| | GLG23619 | southern Hengduan | 2023 | blood | 98.75834 | 24.93019 | 11.6 |
| | GLGS5798* | southern Hengduan | 2005 | muscle | 95.17747 | 29.24492 | 14.9 |
| *Aegithalos concinnus* | 2022LS242 | southern Hengduan | 2022 | muscle | 98.71339 | 25.95805 | 12.2 |
| | 2022LS260 | southern Hengduan | 2022 | muscle | 98.71339 | 25.95805 | 12.9 |
| | GLG23117 | southern Hengduan | 2023 | blood | 98.787 | 25.29984 | 14.8 |
| | GLG23118 | southern Hengduan | 2023 | blood | 98.787 | 25.29984 | 10.7 |
| | GLGS1448* | southern Hengduan | 2003 | muscle | 95.17747 | 29.24492 | 12.3 |
| *Aegithalos iouschistos* | 2022GS130 | southern Hengduan | 2022 | muscle | 98.68341 | 25.97262 | 14.7 |
| | 2022GS131 | southern Hengduan | 2022 | muscle | 98.68341 | 25.97262 | 13.5 |
| | 2022GS132 | southern Hengduan | 2022 | muscle | 98.68341 | 25.97262 | 13.7 |
| | 2022LS107 | southern Hengduan | 2022 | muscle | 98.68341 | 25.97262 | 14.4 |
| | 2022LS141 | southern Hengduan | 2022 | muscle | 98.713 | 25.964 | 17.3 |
| *Aethopyga gouldiae* | 05157 | southern Hengduan | 2005 | muscle | 100.1169 | 24.56964 | 10.4 |
| | GLGS2150 | southern Hengduan | 2004 | muscle | 98.66333 | 25.97294 | 9.5 |
| | GLGS6145 | southern Hengduan | 2006 | muscle | 98.61431 | 25.77742 | 12.3 |
| | GLGS6254 | southern Hengduan | 2006 | muscle | 98.61447 | 25.7658 | 16.6 |
| | GLGS6285 | southern Hengduan | 2006 | muscle | 98.61494 | 25.7658 | 10.4 |
| | 2022LS162 | southern Hengduan | 2022 | muscle | 98.69904 | 25.94157 | 18 |

| | | | | | | | |
|---|---|---|---|---|---|---|---|
| *Aethopyga ignicauda* | 2022LS144 | southern Hengduan | 2022 | muscle | 98.713 | 25.95805 | 15.5 |
| | 2022LS145 | southern Hengduan | 2022 | muscle | 98.713 | 25.95805 | 14.3 |
| | 2022LS146 | southern Hengduan | 2022 | muscle | 98.713 | 25.95805 | 13.8 |
| | 2022LS170 | southern Hengduan | 2022 | muscle | 98.69904 | 25.94157 | 16.4 |
| *Aethopyga nipalensis* | GLGS1365 | southern Hengduan | 2003 | muscle | 98.75892 | 24.85692 | 12.8 |
| | GLGS1410 | southern Hengduan | 2003 | muscle | 98.76631 | 24.82972 | 14.2 |
| | GLGS2129 | southern Hengduan | 2004 | muscle | 98.66333 | 25.99422 | 10.5 |
| | GLGS6313 | southern Hengduan | 2006 | muscle | 98.61767 | 25.80407 | 8.9 |
| | GLGS6314 | southern Hengduan | 2006 | muscle | 98.61767 | 25.80407 | 10.2 |
| *Alcippe castaneceps* | 04154 | southern Hengduan | 2004 | muscle | 98.3329 | 27.92983 | 9.8 |
| | GLGS2102 | southern Hengduan | 2004 | muscle | 98.65925 | 26.00803 | 9.6 |
| | GLGS2134 | southern Hengduan | 2004 | muscle | 98.66333 | 25.99422 | 8.8 |
| | GLGS2223 | southern Hengduan | 2004 | muscle | 98.70508 | 25.98519 | 8.3 |
| | GLGS2360 | southern Hengduan | 2004 | muscle | 98.71306 | 25.97686 | 10.6 |
| *Alcippe chrysotis* | 2022LS014 | southern Hengduan | 2022 | muscle | 98.62972 | 26.02363 | 17 |
| | 2022LS018 | southern Hengduan | 2022 | muscle | 98.62972 | 26.02363 | 17.3 |
| | 2022LS019 | southern Hengduan | 2022 | muscle | 98.62972 | 26.02363 | 12.6 |
| | GLGS2384* | southern Hengduan | 2004 | muscle | 95.9426 | 29.77394 | 19.7 |
| | GLGS5232 | southern Hengduan | 2005 | muscle | 98.71006 | 25.97447 | 15.5 |
| *Alcippe cinerea* | 04043 | southern Hengduan | 2004 | muscle | 98.35997 | 27.89838 | 12.7 |
| | 04044 | southern Hengduan | 2004 | muscle | 98.35997 | 27.89838 | 9.6 |
| | 04053 | southern Hengduan | 2004 | muscle | 98.35997 | 27.89838 | 9.2 |
| | 04057 | southern Hengduan | 2004 | muscle | 98.35997 | 27.89838 | 8.9 |
| | 04058 | southern Hengduan | 2004 | muscle | 98.35997 | 27.89838 | 14.5 |
| *Alcippe dubia* | DHLC041 | southern Hengduan | 2014 | muscle | 97.83313 | 24.72795 | 13.6 |
| | DHYJ016 | southern Hengduan | 2014 | muscle | 97.64067 | 24.63847 | 15.3 |
| | DHYJ020 | southern Hengduan | 2014 | muscle | 97.64067 | 24.63847 | 13.5 |
| | GLG23557 | southern Hengduan | 2023 | blood | 98.82276 | 24.93845 | 11.6 |
| | GLGS1427* | southern Hengduan | 2003 | muscle | 95.6484 | 29.72893 | 12.6 |
| *Alcippe ludlowi* | MT070 | eastern Himalayas | 2018 | blood | 95.6484 | 29.72893 | 15.4 |
| | MT071 | eastern Himalayas | 2018 | blood | 95.6484 | 29.72893 | 18.5 |
| | MT079 | eastern Himalayas | 2018 | blood | 95.64763 | 29.72734 | 22.5 |
| | MT101 | eastern Himalayas | 2018 | blood | 95.64801 | 29.72855 | 17 |

| | | | | | | | |
|---|---|---|---|---|---|---|---|
| | MT069* | eastern Himalayas | 2018 | blood | 95.70076 | 29.8753 | 11.9 |
| *Alcippe morrisonia* | MT420 | eastern Himalayas | 2023 | blood | 95.17699 | 29.24506 | 10 |
| | MT463 | eastern Himalayas | 2023 | blood | 95.17765 | 29.24599 | 9.2 |
| | MT466 | eastern Himalayas | 2023 | blood | 95.17816 | 29.24692 | 12 |
| | MT480 | eastern Himalayas | 2023 | blood | 95.17741 | 29.12477 | 15.5 |
| | MT156* | eastern Himalayas | 2018 | blood | 95.70112 | 29.87534 | 10.9 |
| *Alcippe vinipectus* | GLGS2238 | southern Hengduan | 2004 | muscle | 98.70633 | 25.98514 | 32.8 |
| | GLGS2267 | southern Hengduan | 2004 | muscle | 98.68369 | 25.97331 | 7.7 |
| | GLGS2273 | southern Hengduan | 2004 | muscle | 98.68264 | 25.97197 | 8 |
| | GLGS2373 | southern Hengduan | 2004 | muscle | 98.71044 | 25.98694 | 10.9 |
| | GLGS2376 | southern Hengduan | 2004 | muscle | 98.71086 | 25.98667 | 9.2 |
| *Anthus hodgsoni* | 2022LS050 | southern Hengduan | 2022 | muscle | 98.66696 | 26.05665 | 11.5 |
| | 2022LS084 | southern Hengduan | 2022 | muscle | 98.66696 | 26.05665 | 12.9 |
| | 2022LS094 | southern Hengduan | 2022 | muscle | 98.66696 | 26.05665 | 10.4 |
| | 2022LS103 | southern Hengduan | 2022 | muscle | 98.66696 | 26.05665 | 13.6 |
| | 2022LS102* | southern Hengduan | 2022 | muscle | 95.47852 | 29.49969 | 12.7 |
| *Arachnothera magna* | MT425 | eastern Himalayas | 2023 | muscle | 95.17698 | 29.24691 | 14.1 |
| | MT461 | eastern Himalayas | 2023 | blood | 95.18027 | 29.24741 | 15.3 |
| | MT501 | eastern Himalayas | 2023 | muscle | 95.18277 | 29.24446 | 14.7 |
| | MT527 | eastern Himalayas | 2023 | muscle | 95.18277 | 29.24446 | 12.8 |
| | MT666* | eastern Himalayas | 2020 | blood | 95.17699 | 29.24506 | 12.8 |
| *Carpodacus edwardsii* | 2022LS182 | southern Hengduan | 2022 | muscle | 98.68341 | 25.96893 | 15.1 |
| | 2022LS197 | southern Hengduan | 2022 | muscle | 98.68341 | 25.96893 | 15.1 |
| | GLG23592 | southern Hengduan | 2023 | blood | 98.75797 | 24.93041 | 17 |
| | GLG23613 | southern Hengduan | 2023 | blood | 98.75834 | 24.93018 | 17.2 |
| | GLGS5270 | southern Hengduan | 2005 | muscle | 98.73769 | 27.17722 | 16.9 |
| *Carpodacus erythrinus* | 2022GS074 | southern Hengduan | 2022 | muscle | 98.35072 | 27.69012 | 10.1 |
| | 2022GS083 | southern Hengduan | 2022 | muscle | 98.35065 | 27.69028 | 12.2 |
| | 2022GS088 | southern Hengduan | 2022 | muscle | 98.35047 | 27.69045 | 12.6 |
| | 2022GS092 | southern Hengduan | 2022 | muscle | 98.35072 | 27.69012 | 11.6 |
| | GLGS5108 | southern Hengduan | 2005 | muscle | 98.71042 | 25.98694 | 12.6 |
| *Carpodacus nipalensis* | DHYJ033 | southern Hengduan | 2014 | muscle | 97.69839 | 24.63172 | 10.4 |
| | DHYJ034 | southern Hengduan | 2014 | muscle | 97.69839 | 24.63172 | 12.3 |

| | | | | | | | |
|---|---|---|---|---|---|---|---|
| | DHYJ035 | southern Hengduan | 2014 | muscle | 97.69839 | 24.63172 | 13.3 |
| | GLGS2185 | southern Hengduan | 2004 | muscle | 98.71108 | 25.97792 | 13.8 |
| | GLGS2286 | southern Hengduan | 2004 | muscle | 98.68167 | 25.97294 | 13.7 |
| *Carpodacus thura* | MT049 | eastern Himalayas | 2018 | blood | 95.69757 | 29.80727 | 14.1 |
| | MT050 | eastern Himalayas | 2018 | blood | 95.69757 | 29.80727 | 17.5 |
| | MT053 | eastern Himalayas | 2018 | blood | 95.69757 | 29.80727 | 11.7 |
| | MT343 | eastern Himalayas | 2018 | blood | 95.94032 | 29.77333 | 13.8 |
| | MT005* | eastern Himalayas | 2018 | blood | 95.4551 | 29.47404 | 11.1 |
| *Carpodacus vinaceus* | GLG23093 | southern Hengduan | 2023 | blood | 98.787 | 25.29984 | 14.5 |
| | GLG23094 | southern Hengduan | 2023 | blood | 98.787 | 25.29984 | 18.1 |
| | GLGS1408 | southern Hengduan | 2003 | muscle | 98.76669 | 24.82931 | 14.1 |
| | GLGS5101 | southern Hengduan | 2005 | muscle | 98.71042 | 25.98694 | 15.9 |
| | GLG23158* | southern Hengduan | 2023 | blood | 95.69757 | 29.80727 | 14.6 |
| *Certhia familiaris* | MT317 | eastern Himalayas | 2018 | blood | 95.94086 | 29.77398 | 18.2 |
| | MT322 | eastern Himalayas | 2018 | blood | 95.94189 | 29.77446 | 13.6 |
| | MT335 | eastern Himalayas | 2018 | blood | 95.94032 | 29.77333 | 15.3 |
| | MT020* | eastern Himalayas | 2018 | blood | 95.69757 | 29.80727 | 12 |
| | MT403 | eastern Himalayas | 2020 | blood | 95.33519 | 29.31587 | 12.9 |
| *Cettia flavolivacea* | GLGS2221 | southern Hengduan | 2004 | muscle | 98.70556 | 25.98525 | 14.7 |
| | GLGS2379 | southern Hengduan | 2004 | muscle | 98.71086 | 25.98667 | 12 |
| | GLGS5239 | southern Hengduan | 2005 | muscle | 98.70806 | 27.21164 | 22.5 |
| | GLGS0222 | southern Hengduan | 2002 | muscle | 98.69564 | 25.98525 | 12.9 |
| | GLGS5279* | southern Hengduan | 2005 | muscle | 95.69757 | 29.80727 | 13.5 |
| *Cettia fortipes* | 2022LS240 | southern Hengduan | 2022 | muscle | 98.71339 | 25.95805 | 9 |
| | 2022GS079 | southern Hengduan | 2022 | muscle | 98.35005 | 27.69315 | 10 |
| | 2022GS076 | southern Hengduan | 2022 | muscle | 98.35051 | 27.6952 | 13.6 |
| | 2022LS276 | southern Hengduan | 2022 | muscle | 98.71339 | 25.95805 | 18.6 |
| | GLGS5224 | southern Hengduan | 2005 | muscle | 98.70972 | 25.97617 | 16.3 |
| *Chaimarrornis leucocephalus* | GLGS1393 | southern Hengduan | 2003 | muscle | 98.75967 | 24.85608 | 15.4 |
| | GLGS2113 | southern Hengduan | 2004 | muscle | 98.65925 | 26.00803 | 11.6 |
| | GLGS2356 | southern Hengduan | 2004 | muscle | 98.67897 | 25.97989 | 12.2 |
| | GLGS5074 | southern Hengduan | 2005 | muscle | 98.70486 | 25.98536 | 17.7 |
| | GLGS5127 | southern Hengduan | 2005 | muscle | 98.70486 | 25.98536 | 12.9 |

| | | | | | | | |
|---|---|---|---|---|---|---|---|
| *Chelidorhynx hypoxantha* | 2022GS207 | southern Hengduan | 2022 | muscle | 98.77958 | 24.82578 | 17.2 |
| | 2022LS136 | southern Hengduan | 2022 | muscle | 98.713 | 25.95805 | 15 |
| | GLGS1456 | southern Hengduan | 2003 | muscle | 98.77958 | 24.82578 | 20.3 |
| | GLGS1457 | southern Hengduan | 2003 | muscle | 98.77958 | 24.82578 | 16.1 |
| | GLGS1420* | southern Hengduan | 2003 | muscle | 95.17722 | 29.24925 | 15.3 |
| *Chrysomma sinense* | GLG23247 | southern Hengduan | 2023 | blood | 98.79215 | 24.97281 | 13.6 |
| | GLG23248 | southern Hengduan | 2023 | blood | 98.81215 | 24.97281 | 18.6 |
| | GLG23250 | southern Hengduan | 2023 | blood | 98.80215 | 24.97281 | 14.2 |
| | GLG23683 | southern Hengduan | 2023 | blood | 98.76139 | 25.00429 | 18.3 |
| | GLG23673* | southern Hengduan | 2023 | blood | 95.17894 | 29.24925 | 15 |
| *Cinclidium leucurum* | 2022GS004 | southern Hengduan | 2022 | muscle | 98.31914 | 27.69103 | 14.2 |
| | 2022GS019 | southern Hengduan | 2022 | muscle | 98.32296 | 27.69011 | 11.3 |
| | 2022LS087 | southern Hengduan | 2022 | muscle | 98.85046 | 26.70567 | 12 |
| | DHYJ187 | southern Hengduan | 2014 | muscle | 97.75326 | 24.99136 | 12.4 |
| | GLGS6531 | southern Hengduan | 2006 | muscle | 98.80025 | 25.27732 | 13.1 |
| *Cinclus pallasii* | GLGS1368 | southern Hengduan | 2003 | muscle | 98.75967 | 24.85608 | 15.8 |
| | GLGS1369 | southern Hengduan | 2003 | muscle | 98.75967 | 24.85608 | 19.4 |
| | GLGS1394 | southern Hengduan | 2003 | muscle | 98.75967 | 24.85608 | 19.7 |
| | GLGS1401 | southern Hengduan | 2003 | muscle | 98.75967 | 24.85608 | 14.8 |
| | GLGS5810* | southern Hengduan | 2005 | muscle | 95.17588 | 29.24723 | 12.3 |
| *Culicicapa ceylonensis* | 2022GS003 | southern Hengduan | 2022 | muscle | 98.27914 | 27.69103 | 10 |
| | 2022GS005 | southern Hengduan | 2022 | muscle | 98.27915 | 27.69092 | 13.5 |
| | 2022LS005 | southern Hengduan | 2022 | muscle | 98.69819 | 26.07439 | 13.4 |
| | 2022LS239 | southern Hengduan | 2022 | muscle | 98.75797 | 25.95805 | 13.6 |
| | GLGS6286* | southern Hengduan | 2006 | muscle | 95.17455 | 29.24654 | 13 |
| *Cyornis banyumas* | DHRL001 | southern Hengduan | 2014 | muscle | 97.8911 | 25.04728 | 10.9 |
| | DHRL080 | southern Hengduan | 2014 | muscle | 97.90484 | 25.00877 | 9.5 |
| | DHRL085 | southern Hengduan | 2014 | muscle | 97.92048 | 25.00988 | 12.8 |
| | DHYJ061 | southern Hengduan | 2014 | muscle | 97.60008 | 24.70028 | 13.5 |
| | DHYJ078 | southern Hengduan | 2014 | muscle | 97.60008 | 24.66739 | 10.3 |
| *Cyornis poliogenys* | MT476 | eastern Himalayas | 2023 | muscle | 95.17672 | 29.24811 | 14.6 |
| | MT485 | eastern Himalayas | 2023 | blood | 95.17811 | 29.24603 | 7.2 |
| | MT515 | eastern Himalayas | 2023 | blood | 95.17586 | 29.24616 | 18.3 |

| | | | | | | | |
|---|---|---|---|---|---|---|---|
| | MT655 | eastern Himalayas | 2023 | blood | 95.17641 | 29.24724 | 30.9 |
| | MT448* | eastern Himalayas | 2020 | blood | 95.69849 | 29.806 | 11.1 |
| *Cyornis rubeculoides* | DHYJ178 | southern Hengduan | 2014 | muscle | 97.75326 | 24.99136 | 15.9 |
| | NJDL013 | southern Hengduan | 2017 | muscle | 98.32495 | 27.82627 | 16.4 |
| | NJDL021 | southern Hengduan | 2017 | muscle | 98.29883 | 28.13359 | 16.4 |
| | NJDL138 | southern Hengduan | 2017 | muscle | 98.46598 | 27.84657 | 16.7 |
| | NJDL001* | southern Hengduan | 2017 | muscle | 95.64767 | 29.72748 | 14.7 |
| *Dicrurus leucophaeus* | DHLC060 | southern Hengduan | 2014 | muscle | 98.39216 | 25.36795 | 16.3 |
| | DHRL004 | southern Hengduan | 2014 | muscle | 97.8911 | 25.04728 | 12.2 |
| | DHRL009 | southern Hengduan | 2014 | muscle | 97.8911 | 25.04728 | 10.6 |
| | GLG23301 | southern Hengduan | 2023 | blood | 98.81302 | 24.92035 | 11.6 |
| | 2022GS281* | southern Hengduan | 2022 | muscle | 95.6484 | 29.72893 | 15 |
| *Emberiza elegans* | GLGS1264 | southern Hengduan | 2003 | muscle | 98.61664 | 25.80776 | 12.8 |
| | GLGS1265 | southern Hengduan | 2003 | muscle | 98.61664 | 25.80776 | 13.3 |
| | GLGS1356 | southern Hengduan | 2003 | muscle | 98.75981 | 24.85631 | 14.7 |
| | GLGS1385 | southern Hengduan | 2003 | muscle | 98.75981 | 24.85631 | 11.4 |
| | GLGS1464* | southern Hengduan | 2003 | muscle | 95.69859 | 29.80631 | 13.4 |
| *Enicurus maculatus* | GLGS5142 | southern Hengduan | 2005 | muscle | 98.70558 | 25.98533 | 15 |
| | GLGS5209 | southern Hengduan | 2005 | muscle | 98.71172 | 25.97597 | 16.5 |
| | NJDL004 | southern Hengduan | 2017 | muscle | 98.32495 | 27.82627 | 16.4 |
| | GLGS5800 | southern Hengduan | 2005 | muscle | 98.65953 | 25.00933 | 11.6 |
| | GLGS5185* | southern Hengduan | 2005 | muscle | 95.17319 | 29.24741 | 15.2 |
| *Enicurus scouleri* | 2022GS015 | southern Hengduan | 2022 | muscle | 98.28354 | 27.68932 | 16.3 |
| | 2022GS200 | southern Hengduan | 2022 | muscle | 98.28354 | 27.68932 | 15.5 |
| | 2022GS209 | southern Hengduan | 2022 | muscle | 98.28354 | 27.68932 | 17.2 |
| | 2022GS221 | southern Hengduan | 2022 | muscle | 98.28353 | 27.68932 | 16.2 |
| | GLGS2400* | southern Hengduan | 2004 | muscle | 95.17319 | 29.24741 | 12.5 |
| *Erpornis zantholeuca* | DHLC005 | southern Hengduan | 2014 | muscle | 97.83313 | 24.86795 | 15 |
| | DHLC010 | southern Hengduan | 2014 | muscle | 97.83313 | 24.9568 | 15.9 |
| | DHRL023 | southern Hengduan | 2014 | muscle | 97.81551 | 24.99877 | 15.8 |
| | DHRL065 | southern Hengduan | 2014 | muscle | 97.82355 | 24.99877 | 12.9 |
| | DHLC004* | southern Hengduan | 2014 | muscle | 95.17464 | 29.24826 | 11.7 |
| | 2022LS248 | southern Hengduan | 2022 | muscle | 98.71339 | 25.95805 | 13.7 |

| | | | | | | | |
|---|---|---|---|---|---|---|---|
| *Eumyias thalassinus* | DHLC001 | southern Hengduan | 2014 | muscle | 97.83313 | 24.86795 | 13.9 |
| | DHYJ092 | southern Hengduan | 2014 | muscle | 97.60008 | 24.66739 | 14.1 |
| | GLGS2290 | southern Hengduan | 2004 | muscle | 98.68333 | 25.97292 | 13.6 |
| | GLGS6424* | southern Hengduan | 2006 | muscle | 95.17464 | 29.24826 | 12.7 |
| *Ficedula albicilla* | 2022GS045 | southern Hengduan | 2022 | muscle | 98.54179 | 27.68896 | 11.6 |
| | 2022GS078 | southern Hengduan | 2022 | muscle | 98.35051 | 27.6952 | 10.5 |
| | 2022GS223 | southern Hengduan | 2022 | muscle | 98.31179 | 27.68895 | 12.1 |
| | GLGS0117 | southern Hengduan | 2002 | muscle | 98.72035 | 25.97583 | 11.6 |
| | GLGS2283 | southern Hengduan | 2004 | muscle | 98.68247 | 25.97233 | 11.7 |
| *Ficedula hodgsonii* | GLGS1287 | southern Hengduan | 2003 | muscle | 98.62697 | 25.78578 | 11.1 |
| | GLGS1332 | southern Hengduan | 2003 | muscle | 98.61447 | 25.78553 | 11.6 |
| | GLGS1349 | southern Hengduan | 2003 | muscle | 98.61344 | 25.78542 | 13.7 |
| | GLGS1802 | southern Hengduan | 2003 | muscle | 98.78161 | 25.30342 | 10.6 |
| | GLGS1803* | southern Hengduan | 2003 | muscle | 95.17672 | 29.24811 | 15.4 |
| *Ficedula hyperythra* | 2022LS030 | southern Hengduan | 2022 | muscle | 98.61824 | 26.0414 | 15.7 |
| | 2022LS075 | southern Hengduan | 2022 | muscle | 98.66696 | 26.05665 | 16.9 |
| | 2022LS082 | southern Hengduan | 2022 | muscle | 98.61824 | 26.0414 | 15.9 |
| | 2022LS095 | southern Hengduan | 2022 | muscle | 98.66696 | 26.05665 | 15.9 |
| | 2022LS097 | southern Hengduan | 2022 | muscle | 98.61824 | 26.0414 | 15.7 |
| *Ficedula strophiata* | 2022LS060 | southern Hengduan | 2022 | muscle | 98.66696 | 26.05666 | 12.7 |
| | 2022LS061 | southern Hengduan | 2022 | muscle | 98.66696 | 26.05666 | 11.9 |
| | 2022LS077 | southern Hengduan | 2022 | muscle | 98.66696 | 26.05666 | 14.1 |
| | 2022LS111 | southern Hengduan | 2022 | muscle | 98.68341 | 25.96893 | 14.6 |
| | 2022LS112 | southern Hengduan | 2022 | muscle | 98.68341 | 25.96893 | 12.6 |
| *Ficedula tricolor* | 2022GS110 | southern Hengduan | 2022 | muscle | 98.71044 | 25.98667 | 14 |
| | GLGS2370 | southern Hengduan | 2004 | muscle | 98.71044 | 25.98667 | 12.9 |
| | GLGS5042 | southern Hengduan | 2005 | muscle | 98.69981 | 25.97186 | 15 |
| | GLGS5186 | southern Hengduan | 2005 | muscle | 98.70972 | 25.97617 | 13.7 |
| | GLGS5173* | southern Hengduan | 2005 | muscle | 95.17641 | 29.24724 | 11.8 |
| *Garrulax affinis* | 2022LS232 | southern Hengduan | 2022 | muscle | 98.7168 | 25.95805 | 17.5 |
| | 2022LS233 | southern Hengduan | 2022 | muscle | 98.7168 | 25.95805 | 16.7 |
| | GLGS2250 | southern Hengduan | 2004 | muscle | 98.68253 | 25.97292 | 19.9 |
| | NJDL133 | southern Hengduan | 2017 | muscle | 98.26553 | 27.84717 | 15.8 |

| | | | | | | | |
|---|---|---|---|---|---|---|---|
| | NJDL134 | southern Hengduan | 2017 | muscle | 98.26553 | 27.84717 | 15.1 |
| *Garrulax erythrocephalus* | 2022LS128 | southern Hengduan | 2022 | muscle | 98.68341 | 25.96893 | 13.9 |
| | GLGS5728 | southern Hengduan | 2005 | muscle | 98.65955 | 25.99344 | 15.2 |
| | GLGS5729 | southern Hengduan | 2005 | muscle | 98.65955 | 25.99344 | 17.1 |
| | GLGS5094 | southern Hengduan | 2005 | muscle | 98.71089 | 25.98672 | 23.7 |
| | GLGS5104 | southern Hengduan | 2005 | muscle | 98.71089 | 25.98672 | 14.7 |
| *Garrulax subunicolor* | GLGS0279 | southern Hengduan | 2002 | muscle | 98.71706 | 27.20611 | 15.7 |
| | GLGS5191 | southern Hengduan | 2005 | muscle | 98.71089 | 25.97825 | 14.3 |
| | GLGS5192 | southern Hengduan | 2005 | muscle | 98.71089 | 25.97825 | 16.5 |
| | GLGS5770 | southern Hengduan | 2005 | muscle | 98.65957 | 25.99344 | 17.3 |
| | GLGS2167* | southern Hengduan | 2004 | muscle | 95.47757 | 29.49919 | 13.5 |
| *Hemixos flavala* | MT446 | eastern Himalayas | 2023 | blood | 95.18019 | 29.2486 | 14.9 |
| | MT451 | eastern Himalayas | 2023 | blood | 95.18019 | 29.2486 | 13 |
| | MT535 | eastern Himalayas | 2023 | blood | 95.17665 | 29.2464 | 12.2 |
| | MT540 | eastern Himalayas | 2023 | blood | 95.17665 | 29.2464 | 14.3 |
| | MT541* | eastern Himalayas | 2020 | blood | 95.47757 | 29.49919 | 12 |
| *Heterophasia desgodinsi* | GLG23028 | southern Hengduan | 2023 | blood | 98.78655 | 25.29961 | 15.7 |
| | GLG23029 | southern Hengduan | 2023 | blood | 98.78655 | 25.29961 | 16.5 |
| | GLG23030 | southern Hengduan | 2023 | blood | 98.78655 | 25.29961 | 12.7 |
| | GLG23560 | southern Hengduan | 2023 | blood | 98.82276 | 24.93845 | 10.4 |
| | GLGS6416* | southern Hengduan | 2006 | muscle | 95.45539 | 29.47474 | 13.6 |
| *Heterophasia pulchella* | 2022LS029 | southern Hengduan | 2022 | muscle | 98.75824 | 26.2414 | 11.5 |
| | 2022LS230 | southern Hengduan | 2022 | muscle | 98.7168 | 25.95805 | 14.3 |
| | 2022LS231 | southern Hengduan | 2022 | muscle | 98.7168 | 25.95805 | 15.4 |
| | 2022LS274 | southern Hengduan | 2022 | muscle | 98.71339 | 25.95805 | 15.8 |
| | 04036* | southern Hengduan | 2004 | muscle | 95.17665 | 29.2464 | 11.1 |
| *Hypsipetes leucocephalus* | MT422 | eastern Himalayas | 2023 | blood | 95.17319 | 29.24741 | 11.6 |
| | MT431 | eastern Himalayas | 2023 | blood | 95.17319 | 29.24741 | 15.6 |
| | MT458 | eastern Himalayas | 2023 | blood | 95.17464 | 29.24826 | 14.7 |
| | MT473 | eastern Himalayas | 2023 | blood | 95.17464 | 29.24826 | 10.4 |
| | MT671* | eastern Himalayas | 2020 | blood | 95.18019 | 29.2486 | 11.3 |
| *Hypsipetes mcclellandii* | DHLC002 | southern Hengduan | 2014 | muscle | 97.83216 | 24.76795 | 14.1 |
| | DHLC003 | southern Hengduan | 2014 | muscle | 97.83216 | 24.86795 | 13 |

| | | | | | | | |
|---|---|---|---|---|---|---|---|
| | GLG23543 | southern Hengduan | 2023 | blood | 98.80455 | 24.93577 | 12.6 |
| | GLG23561 | southern Hengduan | 2023 | blood | 98.80455 | 24.93577 | 12.2 |
| | GLG23563 | southern Hengduan | 2023 | blood | 98.80455 | 24.93577 | 12.8 |
| *Leiothrix argentauris* | MT439 | eastern Himalayas | 2023 | blood | 95.33519 | 29.31587 | 13.2 |
| | MT454 | eastern Himalayas | 2023 | blood | 95.17672 | 29.24811 | 19.3 |
| | MT455 | eastern Himalayas | 2023 | blood | 95.17672 | 29.24811 | 19.5 |
| | MT456 | eastern Himalayas | 2023 | blood | 95.17672 | 29.24811 | 16.8 |
| | MT663* | eastern Himalayas | 2023 | blood | 95.17665 | 29.2464 | 14.7 |
| *Leiothrix lutea* | 04015 | southern Hengduan | 2004 | muscle | 98.3572 | 27.89868 | 5.6 |
| | 04076 | southern Hengduan | 2004 | muscle | 98.32232 | 28.07478 | 9.1 |
| | 04081 | southern Hengduan | 2004 | muscle | 98.32442 | 28.07577 | 8.7 |
| | 04084 | southern Hengduan | 2004 | muscle | 98.32631 | 28.07625 | 9.2 |
| | 04172 | southern Hengduan | 2004 | muscle | 98.30314 | 27.68572 | 8 |
| *Lonchura punctulata* | GLG23653 | southern Hengduan | 2023 | blood | 98.76304 | 24.87201 | 11.9 |
| | GLG23660 | southern Hengduan | 2023 | blood | 98.71904 | 24.63201 | 12.9 |
| | GLG23675 | southern Hengduan | 2023 | blood | 98.56304 | 24.78201 | 12 |
| | GLG23676 | southern Hengduan | 2023 | blood | 98.63304 | 24.80201 | 10.1 |
| | GLG23677 | southern Hengduan | 2023 | blood | 98.23304 | 24.97201 | 10.9 |
| *Minla cyanouroptera* | GLGS6485* | southern Hengduan | 2023 | muscle | 95.1775 | 29.24667 | 9.5 |
| | GLG23521 | southern Hengduan | 2023 | blood | 98.80331 | 24.93587 | 8.4 |
| | GLG23288 | southern Hengduan | 2023 | blood | 98.79335 | 25.21985 | 10.1 |
| | GLG23289 | southern Hengduan | 2023 | blood | 98.79335 | 25.21985 | 9.4 |
| | GLG23520 | southern Hengduan | 2023 | blood | 98.80331 | 24.93587 | 9.5 |
| *Minla ignotincta* | GLG23522 | southern Hengduan | 2023 | blood | 98.80331 | 24.93587 | 14.5 |
| | GLG23553 | southern Hengduan | 2023 | blood | 98.80331 | 24.93587 | 15.9 |
| | GLGS5793 | southern Hengduan | 2005 | muscle | 98.65967 | 25.00897 | 15.1 |
| | GLGS5794 | southern Hengduan | 2005 | muscle | 98.65967 | 25.00897 | 13.7 |
| | 04469* | southern Hengduan | 2004 | muscle | 95.17808 | 29.24648 | 10.4 |
| *Minla strigula* | 2022GS180 | southern Hengduan | 2022 | muscle | 98.68341 | 25.96893 | 15.7 |
| | 2022GS181 | southern Hengduan | 2022 | muscle | 98.6834 | 25.96894 | 16.9 |
| | 2022LS123 | southern Hengduan | 2022 | muscle | 98.68341 | 25.96893 | 13.3 |
| | 2022LS164 | southern Hengduan | 2022 | muscle | 98.713 | 25.95805 | 10.7 |
| | GLGS5100* | southern Hengduan | 2005 | muscle | 95.17539 | 29.24321 | 14.9 |

| | | | | | | | |
|---|---|---|---|---|---|---|---|
| *Myzornis pyrrhoura* | 2022GS140 | southern Hengduan | 2022 | muscle | 98.69217 | 25.96893 | 29.5 |
| | 2022GS141 | southern Hengduan | 2022 | muscle | 98.68017 | 25.96893 | 12.6 |
| | 2022LS201 | southern Hengduan | 2022 | muscle | 98.61168 | 25.68439 | 14.8 |
| | 2022LS204 | southern Hengduan | 2022 | muscle | 98.79502 | 24.96893 | 17.6 |
| | 2022LS205 | southern Hengduan | 2022 | muscle | 98.74502 | 25.96893 | 17.4 |
| *Niltava grandis* | 2022GS014 | southern Hengduan | 2022 | muscle | 98.27915 | 27.69091 | 11.6 |
| | 2022GS016 | southern Hengduan | 2022 | muscle | 98.28354 | 27.68932 | 11.9 |
| | 2022GS056 | southern Hengduan | 2022 | muscle | 98.31179 | 27.68896 | 17 |
| | 2022GS084 | southern Hengduan | 2022 | muscle | 98.35065 | 27.69028 | 13.2 |
| | GLGS1463* | southern Hengduan | 2003 | muscle | 95.17893 | 29.14447 | 14 |
| *Niltava macgrigoriae* | 2022GS017 | southern Hengduan | 2022 | muscle | 98.28296 | 27.69011 | 15.1 |
| | 2022GS023 | southern Hengduan | 2022 | muscle | 98.28353 | 27.68932 | 15 |
| | 2022GS037 | southern Hengduan | 2022 | muscle | 98.27915 | 27.69091 | 14.7 |
| | 2022GS041 | southern Hengduan | 2022 | muscle | 98.31062 | 27.68872 | 21.8 |
| | 2022LS254 | southern Hengduan | 2022 | muscle | 98.71339 | 25.95805 | 15.5 |
| *Niltava sundara* | 2022GS094 | southern Hengduan | 2022 | muscle | 98.79117 | 26.35805 | 13.3 |
| | 2022GS095 | southern Hengduan | 2022 | muscle | 98.79217 | 25.25805 | 14.1 |
| | 2022LS247 | southern Hengduan | 2022 | muscle | 98.76022 | 26.78958 | 16.9 |
| | 2022LS252 | southern Hengduan | 2022 | muscle | 98.80217 | 25.85758 | 13.5 |
| | 2022LS255 | southern Hengduan | 2022 | muscle | 98.80621 | 25.12036 | 15.4 |
| *Orthotomus sutorius* | GLG23225 | southern Hengduan | 2023 | blood | 98.75834 | 24.93019 | 13.9 |
| | GLG23256 | southern Hengduan | 2023 | blood | 98.73218 | 24.97264 | 14.4 |
| | GLG23267 | southern Hengduan | 2023 | blood | 98.69322 | 24.97281 | 15.8 |
| | GLG23269 | southern Hengduan | 2023 | blood | 98.71322 | 24.97281 | 16.8 |
| | GLG23270 | southern Hengduan | 2023 | blood | 98.71132 | 24.97281 | 15.3 |
| *Paradoxornis fulvifrons* | 2022LS185 | southern Hengduan | 2022 | muscle | 98.68341 | 25.96893 | 16.2 |
| | 2022LS191 | southern Hengduan | 2022 | muscle | 98.68341 | 25.96893 | 13.8 |
| | 2022LS137 | southern Hengduan | 2022 | muscle | 98.713 | 25.95805 | 15 |
| | 2022LS138 | southern Hengduan | 2022 | muscle | 98.713 | 25.95805 | 12.9 |
| | GLGS0352* | southern Hengduan | 2002 | muscle | 95.44713 | 29.48116 | 14.7 |
| *Paradoxornis nipalensis* | GLGS1422 | southern Hengduan | 2003 | muscle | 98.77919 | 24.81967 | 13.1 |
| | GLGS5776 | southern Hengduan | 2005 | muscle | 98.65831 | 25.99464 | 14.5 |

| | | | | | | | |
|---|---|---|---|---|---|---|---|
| | GLGS5777 | southern Hengduan | 2005 | muscle | 98.65831 | 25.99464 | 13.3 |
| | GLGS5778 | southern Hengduan | 2005 | muscle | 98.65831 | 25.99464 | 12.9 |
| | GLGS5775 | southern Hengduan | 2005 | muscle | 98.65831 | 25.99464 | 13.7 |
| *Parus ater* | MT098 | eastern Himalayas | 2018 | blood | 95.64802 | 29.72855 | 10.8 |
| | MT104 | eastern Himalayas | 2018 | blood | 95.64802 | 29.72855 | 22.9 |
| | MT417 | eastern Himalayas | 2023 | blood | 95.47757 | 29.49919 | 12.7 |
| | MT419 | eastern Himalayas | 2023 | muscle | 95.47757 | 29.49919 | 16.8 |
| | MT086 | eastern Himalayas | 2018 | blood | 95.6484 | 29.72893 | 13.2 |
| *Parus monticolus* | 2022GS184 | southern Hengduan | 2022 | muscle | 98.7114 | 25.96893 | 11 |
| | 2022LS195 | southern Hengduan | 2022 | muscle | 98.71398 | 25.96893 | 15.7 |
| | GLG23234 | southern Hengduan | 2023 | blood | 98.75834 | 24.93019 | 13.8 |
| | GLG23312 | southern Hengduan | 2023 | blood | 98.79378 | 25.21946 | 14.8 |
| | GLG23608 | southern Hengduan | 2023 | blood | 98.75777 | 24.92989 | 17.3 |
| *Parus rubidiventris* | 2022LS167 | southern Hengduan | 2022 | muscle | 98.713 | 25.95805 | 18.7 |
| | 2022LS200 | southern Hengduan | 2022 | muscle | 98.70411 | 25.96893 | 19.5 |
| | 2022LS203 | southern Hengduan | 2022 | muscle | 98.70041 | 25.96893 | 13.4 |
| | GLGS0220 | southern Hengduan | 2002 | muscle | 98.71886 | 27.21178 | 15.8 |
| | GLGS5240 | southern Hengduan | 2005 | muscle | 98.71919 | 27.21053 | 16.8 |
| *Parus spilonotus* | GLG23131 | southern Hengduan | 2023 | blood | 98.78757 | 25.30038 | 14.3 |
| | GLG23170 | southern Hengduan | 2023 | blood | 98.8025 | 25.29802 | 11.6 |
| | NJDL002 | southern Hengduan | 2017 | muscle | 98.32495 | 27.82627 | 13.9 |
| | NJDL003 | southern Hengduan | 2017 | muscle | 98.32495 | 27.82627 | 15.4 |
| | NJDL012 | southern Hengduan | 2017 | muscle | 98.32495 | 27.82627 | 13.9 |
| *Passer rutilans* | MT166 | eastern Himalayas | 2018 | blood | 95.454 | 29.47356 | 16.3 |
| | MT168 | eastern Himalayas | 2018 | blood | 95.454 | 29.47356 | 15.7 |
| | MT171 | eastern Himalayas | 2018 | blood | 95.454 | 29.47356 | 15.8 |
| | MT172 | eastern Himalayas | 2018 | blood | 95.454 | 29.47356 | 16 |
| | MT170* | eastern Himalayas | 2018 | blood | 95.6484 | 29.72893 | 11.3 |
| *Phoenicurus frontalis* | 2022LS078 | southern Hengduan | 2022 | muscle | 98.66696 | 26.05666 | 13.9 |
| | 2022LS163 | southern Hengduan | 2022 | muscle | 98.713 | 25.95805 | 12.3 |
| | 2022LS166 | southern Hengduan | 2022 | muscle | 98.713 | 25.95805 | 13.9 |
| | GLG23310 | southern Hengduan | 2023 | blood | 98.79335 | 25.21985 | 19.8 |
| | GLG23616 | southern Hengduan | 2023 | blood | 98.75834 | 24.93019 | 9.5 |

| | | | | | | | |
|---|---|---|---|---|---|---|---|
| *Phoenicurus schisticeps* | MT314 | eastern Himalayas | 2018 | blood | 95.94088 | 29.7737 | 12.2 |
| | MT320 | eastern Himalayas | 2018 | blood | 95.94155 | 29.77433 | 15.4 |
| | MT323 | eastern Himalayas | 2018 | blood | 95.94088 | 29.7737 | 12.6 |
| | MT324 | eastern Himalayas | 2018 | blood | 95.9426 | 29.77394 | 19.1 |
| | MT313* | eastern Himalayas | 2018 | blood | 95.64802 | 29.72855 | 12.5 |
| *Phylloscopus affinis* | GLGS5008 | southern Hengduan | 2005 | muscle | 98.68258 | 25.97258 | 11.5 |
| | GLGS5031 | southern Hengduan | 2005 | muscle | 98.68366 | 25.97239 | 13 |
| | GLGS5202 | southern Hengduan | 2005 | muscle | 98.70972 | 25.97617 | 12 |
| | GLGS5222 | southern Hengduan | 2005 | muscle | 98.70964 | 25.97639 | 12.6 |
| | GLGS5021 | southern Hengduan | 2005 | muscle | 98.68372 | 25.97322 | 19.1 |
| *Phylloscopus armandii* | GLG23211 | southern Hengduan | 2023 | blood | 98.82333 | 24.93797 | 7.8 |
| | GLG23213 | southern Hengduan | 2023 | blood | 98.82333 | 24.93797 | 14.3 |
| | GLGS0432 | southern Hengduan | 2002 | muscle | 98.71822 | 27.20661 | 15.9 |
| | GLGS5115 | southern Hengduan | 2005 | muscle | 98.71089 | 25.98672 | 14.1 |
| | 2022GS206* | southern Hengduan | 2022 | muscle | 95.47757 | 29.49919 | 11 |
| *Phylloscopus cantator* | MT511 | eastern Himalayas | 2023 | blood | 95.1775 | 29.24667 | 16.2 |
| | MT537 | eastern Himalayas | 2023 | muscle | 95.17808 | 29.24648 | 17.6 |
| | MT561 | eastern Himalayas | 2023 | blood | 95.17539 | 29.24321 | 11.6 |
| | MT578 | eastern Himalayas | 2023 | blood | 95.17905 | 29.24533 | 12.3 |
| | MT530* | eastern Himalayas | 2020 | blood | 95.6484 | 29.72893 | 11.4 |
| *Phylloscopus chloronotus* | MT011 | eastern Himalayas | 2018 | blood | 95.69849 | 29.806 | 8.7 |
| | MT030 | eastern Himalayas | 2018 | blood | 95.69849 | 29.806 | 10.5 |
| | MT064 | eastern Himalayas | 2018 | blood | 95.64767 | 29.72748 | 13.4 |
| | MT078 | eastern Himalayas | 2018 | blood | 95.6484 | 29.72893 | 13.4 |
| | MT002* | eastern Himalayas | 2018 | blood | 95.454 | 29.47356 | 9 |
| *Phylloscopus davisoni* | DHLC009 | southern Hengduan | 2014 | muscle | 97.83313 | 24.95795 | 12.5 |
| | DHYJ203 | southern Hengduan | 2014 | muscle | 97.75326 | 24.99136 | 13.1 |
| | GLGS1323 | southern Hengduan | 2003 | muscle | 98.62719 | 25.78592 | 11.8 |
| | GLGS1397 | southern Hengduan | 2003 | muscle | 98.76033 | 24.85558 | 13.3 |
| | 2022GS048* | southern Hengduan | 2022 | muscle | 95.454 | 29.47356 | 12.4 |
| *Phylloscopus fuscatus* | GLG23194 | southern Hengduan | 2023 | blood | 98.75843 | 25.29055 | 10.5 |
| | GLG23252 | southern Hengduan | 2002 | blood | 98.79817 | 24.97366 | 11.2 |
| | GLGS0080 | southern Hengduan | 2002 | muscle | 98.87425 | 25.70325 | 13.6 |

| | | | | | | | |
|---|---|---|---|---|---|---|---|
| | GLGS0082 | southern Hengduan | 2002 | muscle | 98.87556 | 25.7035 | 10.9 |
| | GLGS0085 | southern Hengduan | 2002 | blood | 98.87467 | 25.70406 | 8.5 |
| *Phylloscopu s maculipenni s* | 2022LS096 | southern Hengduan | 2022 | muscle | 98.61824 | 26.0414 | 19.3 |
| | 2022LS098 | southern Hengduan | 2022 | muscle | 98.61824 | 26.0414 | 11.3 |
| | 2022LS134 | southern Hengduan | 2022 | muscle | 98.7168 | 25.95805 | 15.3 |
| | 2022LS228 | southern Hengduan | 2022 | muscle | 98.7168 | 25.95805 | 13.8 |
| | 04190* | southern Hengduan | 2004 | muscle | 95.454 | 29.47356 | 11.7 |
| *Phylloscopu s magnirostri s* | GLGS2253 | southern Hengduan | 2004 | muscle | 98.67897 | 25.97989 | 14.8 |
| | GLGS2255 | southern Hengduan | 2004 | muscle | 98.67897 | 25.97989 | 13.1 |
| | GLGS2298 | southern Hengduan | 2004 | muscle | 98.67844 | 25.97981 | 17.1 |
| | GLGS2380 | southern Hengduan | 2004 | muscle | 98.70856 | 25.98547 | 17.9 |
| | GLGS6479 | southern Hengduan | 2006 | muscle | 98.84798 | 24.84132 | 14.5 |
| *Phylloscopu s pulcher* | 2022GS099 | southern Hengduan | 2022 | muscle | 98.75696 | 26.05666 | 11.7 |
| | 2022LS076 | southern Hengduan | 2022 | muscle | 98.71696 | 26.05666 | 21.8 |
| | 2022LS099 | southern Hengduan | 2022 | muscle | 98.88043 | 26.7014 | 11.4 |
| | GLG23001 | southern Hengduan | 2023 | blood | 98.787 | 25.29984 | 14.6 |
| | GLG23009 | southern Hengduan | 2023 | blood | 98.787 | 25.29984 | 11.2 |
| *Phylloscopu s xanthoschist os* | MT624 | eastern Himalayas | 2023 | blood | 95.4829 | 29.50108 | 10.8 |
| | MT637 | eastern Himalayas | 2023 | blood | 95.47757 | 29.49919 | 11.5 |
| | MT638 | eastern Himalayas | 2023 | muscle | 95.47757 | 29.49919 | 18 |
| | MT133* | eastern Himalayas | 2018 | blood | 95.18027 | 29.24741 | 11.6 |
| | MT545 | eastern Himalayas | 2020 | blood | 95.17665 | 29.2464 | 12.5 |
| *Pnoepyga pusilla* | 2022LS027 | southern Hengduan | 2022 | muscle | 98.62972 | 26.02363 | 16.4 |
| | GLGS2133 | southern Hengduan | 2004 | muscle | 98.66333 | 25.99422 | 14.1 |
| | GLGS2199 | southern Hengduan | 2004 | muscle | 98.71108 | 25.97703 | 14.7 |
| | GLGS6153 | southern Hengduan | 2006 | muscle | 98.61847 | 25.77976 | 17.3 |
| | GLGS6346 | southern Hengduan | 2006 | muscle | 98.6982 | 25.65651 | 20.4 |
| *Pomatorhin us ruficollis* | GLG23222 | southern Hengduan | 2023 | blood | 98.82285 | 24.93848 | 12.1 |
| | GLG23287 | southern Hengduan | 2023 | blood | 98.79378 | 25.21946 | 16.6 |
| | GLG23294 | southern Hengduan | 2023 | blood | 98.79268 | 25.21935 | 18.7 |
| | GLG23641 | southern Hengduan | 2023 | blood | 98.75834 | 24.93018 | 13.4 |
| | GLG23135* | southern Hengduan | 2023 | blood | 95.18277 | 29.24446 | 21.9 |
| | 2022GS075 | southern Hengduan | 2022 | muscle | 98.35065 | 27.69028 | 12.8 |

| | | | | | | | |
|---|---|---|---|---|---|---|---|
| *Prinia atrogularis* | 2022GS077 | southern Hengduan | 2022 | muscle | 98.35051 | 27.6952 | 13.5 |
| | GLG23148 | southern Hengduan | 2023 | blood | 98.78677 | 25.30017 | 20.3 |
| | GLG23215 | southern Hengduan | 2023 | blood | 98.82221 | 24.93822 | 12.6 |
| | 04427 | southern Hengduan | 2004 | muscle | 98.35178 | 27.74275 | 12 |
| *Prinia hodgsonii* | GLG23207 | southern Hengduan | 2023 | blood | 98.8223 | 24.93814 | 15.4 |
| | GLG23208 | southern Hengduan | 2023 | blood | 98.8223 | 24.93814 | 14.9 |
| | GLG23219 | southern Hengduan | 2023 | blood | 98.82276 | 24.93845 | 14.3 |
| | GLG23515 | southern Hengduan | 2023 | blood | 98.80461 | 25.2846 | 29 |
| | GLG23691* | southern Hengduan | 2023 | blood | 95.94086 | 29.77398 | 14.4 |
| *Prunella immaculata* | 2022LS058 | southern Hengduan | 2022 | muscle | 98.66696 | 26.05665 | 14.4 |
| | 2022LS059 | southern Hengduan | 2022 | muscle | 98.66696 | 26.05665 | 13.2 |
| | 2022LS085 | southern Hengduan | 2022 | muscle | 98.66696 | 26.05665 | 18.8 |
| | 2022LS086 | southern Hengduan | 2022 | muscle | 98.66696 | 26.05665 | 14.9 |
| | 2022LS198 | southern Hengduan | 2022 | muscle | 98.68341 | 25.96893 | 11.4 |
| *Prunella* strophiata | MT212 | eastern Himalayas | 2018 | blood | 95.70018 | 29.87567 | 15.7 |
| | MT218 | eastern Himalayas | 2018 | blood | 95.70075 | 29.87569 | 13.7 |
| | MT230 | eastern Himalayas | 2018 | blood | 95.69992 | 29.87557 | 14 |
| | MT242 | eastern Himalayas | 2018 | blood | 95.70023 | 29.87602 | 15.2 |
| | MT177* | eastern Himalayas | 2018 | blood | 95.94032 | 29.77333 | 10.8 |
| *Pteruthius melanotis* | 2022GS287 | southern Hengduan | 2022 | muscle | 98.61131 | 27.63111 | 16.3 |
| | DHYJ049 | southern Hengduan | 2014 | muscle | 97.67558 | 24.63339 | 17.4 |
| | NJDL011 | southern Hengduan | 2017 | muscle | 98.32495 | 27.82627 | 17.7 |
| | GLGS6268 | southern Hengduan | 2006 | muscle | 98.6982 | 25.7658 | 16.6 |
| | 04150* | southern Hengduan | 2004 | muscle | 95.69757 | 29.80727 | 14.9 |
| *Pycnonotus cafer* | GLG23261 | southern Hengduan | 2023 | blood | 98.54866 | 24.97366 | 14.9 |
| | GLG23282 | southern Hengduan | 2023 | blood | 98.54866 | 24.97281 | 13.7 |
| | GLG23283 | southern Hengduan | 2023 | blood | 98.54866 | 24.97281 | 13.2 |
| | GLG23692 | southern Hengduan | 2023 | blood | 98.54866 | 25.00319 | 12.6 |
| | GLG23685 | southern Hengduan | 2023 | blood | 98.54866 | 24.97201 | 13.2 |
| *Pycnonotus flavescens* | DHYJ054 | southern Hengduan | 2014 | muscle | 97.63172 | 24.63172 | 11.6 |
| | DHYJ116 | southern Hengduan | 2014- | muscle | 97.75326 | 24.99136 | 13.4 |
| | DHYJ162 | southern Hengduan | 2014 | muscle | 97.75326 | 24.99136 | 12.5 |
| | DHYJ177 | southern Hengduan | 2014 | muscle | 97.75326 | 24.99136 | 13.2 |

| | | | | | | | |
|---|---|---|---|---|---|---|---|
| | GLG23239* | southern Hengduan | 2023 | blood | 95.33519 | 29.31587 | 15.4 |
| *Pycnonotus xanthorrhous* | 2022GS085 | southern Hengduan | 2022 | muscle | 98.35003 | 27.693 | 15 |
| | GLG23206 | southern Hengduan | 2023 | blood | 98.82276 | 24.93845 | 11.5 |
| | GLG23231 | southern Hengduan | 2023 | blood | 98.75777 | 24.92989 | 25.3 |
| | GLG23232 | southern Hengduan | 2023 | blood | 98.75777 | 24.92989 | 16.7 |
| | YL20080017* | southern Hengduan | 2008 | muscle | 95.17672 | 29.24811 | 14.9 |
| *Pyrrhoplectes epauletta* | GLGS0267 | southern Hengduan | 2002 | muscle | 98.71108 | 27.21064 | 16.9 |
| | GLGS1484 | southern Hengduan | 2003 | muscle | 98.76703 | 25.30944 | 14.5 |
| | GLGS5064 | southern Hengduan | 2005 | muscle | 98.68226 | 25.97281 | 18.8 |
| | GLGS5088 | southern Hengduan | 2005 | muscle | 98.71031 | 25.98772 | 16.9 |
| | GLGS6315 | southern Hengduan | 2006 | muscle | 98.50016 | 25.80407 | 16.2 |
| *Rhipidura albicollis* | 2022GS035 | southern Hengduan | 2022 | muscle | 98.2832 | 27.68982 | 15 |
| | 2022GS227 | southern Hengduan | 2022 | muscle | 98.2832 | 27.68982 | 17.9 |
| | 2022GS266 | southern Hengduan | 2022 | muscle | 98.30131 | 27.70011 | 17.7 |
| | 2022LS007 | southern Hengduan | 2022 | muscle | 98.60511 | 26.07439 | 15.5 |
| | 2022LS016 | southern Hengduan | 2022 | muscle | 98.62972 | 26.02363 | 12.1 |
| *Rhyacornis fuliginosa* | 2022GS222 | southern Hengduan | 2022 | muscle | 98.33809 | 25.13578 | 13 |
| | GLGS1492 | southern Hengduan | 2003 | muscle | 98.76606 | 25.31261 | 16.7 |
| | GLGS2063 | southern Hengduan | 2004 | muscle | 98.61983 | 26.06225 | 18.2 |
| | GLGS2066 | southern Hengduan | 2004 | muscle | 98.61983 | 26.06225 | 17.9 |
| | 04423* | southern Hengduan | 2004 | muscle | 95.66562 | 29.43575 | 11.6 |
| *Saxicola ferreus* | MT198* | eastern Himalayas | 2018 | blood | 95.70018 | 29.87567 | 14.8 |
| | GLG23204 | southern Hengduan | 2023 | blood | 98.823 | 24.93838 | 16.1 |
| | GLG23205 | southern Hengduan | 2023 | blood | 98.823 | 24.93838 | 14.1 |
| | GLG23214 | southern Hengduan | 2023 | blood | 98.8223 | 24.93814 | 18 |
| | GLGS1838 | southern Hengduan | 2003 | muscle | 98.80325 | 25.29547 | 18.1 |
| *Seicercus burkii* | MT125 | eastern Himalayas | 2018 | blood | 95.45462 | 29.47416 | 18.2 |
| | MT144 | eastern Himalayas | 2018 | blood | 95.45462 | 29.47416 | 13.1 |
| | MT409 | eastern Himalayas | 2023 | blood | 95.44713 | 29.48116 | 12.3 |
| | MT410 | eastern Himalayas | 2023 | blood | 95.44713 | 29.48116 | 9.6 |
| | MT062* | eastern Himalayas | 2018 | blood | 95.70075 | 29.87569 | 10 |
| *Seicercus castaniceps* | 2022LS001 | southern Hengduan | 2022 | muscle | 98.60511 | 26.07439 | 17.1 |
| | 2022LS063 | southern Hengduan | 2022 | muscle | 98.61824 | 26.0414 | 15.4 |

| | | | | | | | |
|---|---|---|---|---|---|---|---|
| | 2022LS214 | southern Hengduan | 2022 | muscle | 98.7168 | 25.95805 | 16.1 |
| | 2022LS225 | southern Hengduan | 2022 | muscle | 98.7168 | 25.95805 | 15 |
| | GLGS2155* | southern Hengduan | 2004 | muscle | 95.69992 | 29.87557 | 13.9 |
| *Seicercus poliogenys* | 2022LS003 | southern Hengduan | 2022 | muscle | 98.60511 | 26.07439 | 16.6 |
| | 2022LS004 | southern Hengduan | 2022 | muscle | 98.60511 | 26.07439 | 12.9 |
| | 2022LS011 | southern Hengduan | 2022 | muscle | 98.60511 | 26.07439 | 17.2 |
| | 2022LS023 | southern Hengduan | 2022 | muscle | 98.60511 | 26.07439 | 15.2 |
| | 04285* | southern Hengduan | 2004 | muscle | 95.70023 | 29.87602 | 14.2 |
| *Spizixos canifrons* | DHYJ210 | southern Hengduan | 2014 | muscle | 97.75326 | 24.99136 | 11.2 |
| | DHYJ214 | southern Hengduan | 2014 | muscle | 97.75326 | 24.99136 | 8 |
| | GLGS6291 | southern Hengduan | 2006 | muscle | 98.69946 | 24.48264 | 10.4 |
| | GLGS6363 | southern Hengduan | 2006 | muscle | 98.69946 | 24.48264 | 9.7 |
| | GLGS6462* | southern Hengduan | 2006 | muscle | 95.45462 | 29.47416 | 11 |
| *Stachyris chrysaea* | 2022GS277 | southern Hengduan | 2022 | muscle | 98.59558 | 27.66591 | 15.3 |
| | 2022GS310 | southern Hengduan | 2022 | muscle | 98.68466 | 27.66657 | 8.3 |
| | 2022GS311 | southern Hengduan | 2022 | muscle | 98.66566 | 27.66561 | 15.2 |
| | NJDL085 | southern Hengduan | 2017 | muscle | 98.71566 | 27.66661 | 10.2 |
| | NJDL112 | southern Hengduan | 2017 | muscle | 98.74553 | 27.66651 | 11.4 |
| *Stachyris nigriceps* | GLGS1854 | southern Hengduan | 2003 | muscle | 98.75817 | 24.57978 | 14.3 |
| | NJDL101 | southern Hengduan | 2017 | muscle | 98.66566 | 27.66661 | 23.1 |
| | NJDL106 | southern Hengduan | 2017 | muscle | 98.69657 | 27.66651 | 17.5 |
| | NJDL109 | southern Hengduan | 2017 | muscle | 98.70657 | 27.66651 | 15.1 |
| | NJDL118 | southern Hengduan | 2017 | muscle | 98.72657 | 27.66651 | 12.8 |
| *Stachyris ruficeps* | 2022LS064 | southern Hengduan | 2022 | muscle | 98.61824 | 26.0414 | 14.5 |
| | 2022LS067 | southern Hengduan | 2022 | muscle | 98.61824 | 26.0414 | 13.5 |
| | 2022LS081 | southern Hengduan | 2022 | muscle | 98.61824 | 26.0414 | 16.2 |
| | 2022LS269 | southern Hengduan | 2022 | muscle | 98.71339 | 25.95805 | 16.2 |
| | GLGS1279 | southern Hengduan | 2003 | muscle | 98.62697 | 25.78578 | 14 |
| *Sylviparus modestus* | 2022GS185 | southern Hengduan | 2022 | muscle | 98.7168 | 25.95805 | 14 |
| | 2022GS186 | southern Hengduan | 2022 | muscle | 98.7168 | 25.95805 | 14 |
| | 2022LS213 | southern Hengduan | 2022 | muscle | 98.7168 | 25.95805 | 12.8 |
| | 2022LS219 | southern Hengduan | 2022 | muscle | 98.7168 | 25.95805 | 12.2 |
| | GLGS2190* | southern Hengduan | 2004 | muscle | 98.71108 | 25.97792 | 13.2 |

| | | | | | | | |
|---|---|---|---|---|---|---|---|
| *Tarsiger chrysaeus* | GLGS1239 | southern Hengduan | 2003 | muscle | 98.62133 | 25.81003 | 11.1 |
| | GLGS1375 | southern Hengduan | 2003 | muscle | 98.76631 | 24.82972 | 9.8 |
| | GLGS1376 | southern Hengduan | 2003 | muscle | 98.76631 | 24.82972 | 10.5 |
| | GLGS2355 | southern Hengduan | 2004 | muscle | 98.68247 | 25.97233 | 9.4 |
| | GLGS2368 | southern Hengduan | 2004 | muscle | 98.71044 | 25.98667 | 7.7 |
| *Tarsiger indicus* | GLGS1238 | southern Hengduan | 2003 | muscle | 98.61756 | 25.80742 | 12.1 |
| | GLGS5011 | southern Hengduan | 2005 | muscle | 98.68242 | 25.97233 | 11.7 |
| | GLGS5040 | southern Hengduan | 2005 | muscle | 98.68255 | 25.97189 | 12.3 |
| | GLGS5243 | southern Hengduan | 2005 | muscle | 98.71272 | 27.20956 | 9.3 |
| | GLGS5726 | southern Hengduan | 2005 | muscle | 98.6595 | 25.99344 | 8.4 |
| *Tephrodorn is gularis* | DHRL006 | southern Hengduan | 2014 | muscle | 97.8911 | 24.68728 | 12.4 |
| | DHRL007 | southern Hengduan | 2014 | muscle | 97.8911 | 24.66728 | 10.6 |
| | DHRL010 | southern Hengduan | 2014 | muscle | 97.8911 | 24.70728 | 13.2 |
| | DHRL082 | southern Hengduan | 2014 | muscle | 97.79551 | 24.65788 | 15 |
| | DHRL011* | southern Hengduan | 2014 | muscle | 97.75326 | 24.99136 | 17.1 |
| *Tesia castaneocor onata* | 2022LS020 | southern Hengduan | 2022 | muscle | 98.62972 | 26.02363 | 13.4 |
| | 2022LS036 | southern Hengduan | 2022 | muscle | 98.61824 | 26.0414 | 14.4 |
| | 2022LS220 | southern Hengduan | 2022 | muscle | 98.7168 | 25.95805 | 23.8 |
| | 2022LS222 | southern Hengduan | 2022 | muscle | 98.7168 | 25.95805 | 15.9 |
| | 2022LS227 | southern Hengduan | 2022 | muscle | 98.7168 | 25.95805 | 17.3 |
| *Tesia cyaniventer* | 2022LS002 | southern Hengduan | 2022 | muscle | 98.60511 | 26.07439 | 16.6 |
| | 2022LS246 | southern Hengduan | 2022 | muscle | 98.71339 | 25.95805 | 15.8 |
| | 2022LS073 | southern Hengduan | 2022 | muscle | 98.61824 | 26.0414 | 12.9 |
| | 2022LS091 | southern Hengduan | 2022 | muscle | 98.61824 | 26.0414 | 16.4 |
| | 04130* | southern Hengduan | 2004 | muscle | 98.76033 | 24.85558 | 11.4 |
| *Turdus albocinctus* | MT100 | eastern Himalayas | 2018 | blood | 95.6484 | 29.72893 | 14.8 |
| | MT211 | eastern Himalayas | 2018 | blood | 95.70076 | 29.8753 | 13.2 |
| | MT227 | eastern Himalayas | 2018 | blood | 95.70112 | 29.87534 | 11.1 |
| | MT072* | eastern Himalayas | 2018 | blood | 98.31179 | 27.68896 | 11.5 |
| | MT408 | eastern Himalayas | 2023 | blood | 95.47852 | 29.49969 | 12 |
| *Turdus dissimilis* | GLGS1348 | southern Hengduan | 2003 | muscle | 98.61494 | 25.78583 | 5.7 |
| | GLGS1498 | southern Hengduan | 2003 | muscle | 98.78161 | 25.30342 | 7.2 |
| | GLGS1499 | southern Hengduan | 2003 | muscle | 98.78161 | 25.30342 | 8.5 |

| | | | | | | | |
|---|---|---|---|---|---|---|---|
| | GLGS1500 | southern Hengduan | 2003 | muscle | 98.78161 | 25.30342 | 7 |
| | DHYJ058* | southern Hengduan | 2003 | muscle | 98.70558 | 25.98533 | 12.2 |
| *Yuhina bakeri* | MT481 | eastern Himalayas | 2023 | blood | 95.17722 | 29.24925 | 12.4 |
| | MT486 | eastern Himalayas | 2023 | blood | 95.17894 | 29.24925 | 12.3 |
| | MT488 | eastern Himalayas | 2023 | blood | 95.24837 | 29.24837 | 13.6 |
| | MT613 | eastern Himalayas | 2023 | blood | 95.17588 | 29.24723 | 12.6 |
| | MT635* | eastern Himalayas | 2020 | blood | 98.71172 | 25.97597 | 11.7 |
| *Yuhina castaniceps* | DHYJ070 | southern Hengduan | 2014 | muscle | 97.56853 | 24.75031 | 11.3 |
| | DHYJ128 | southern Hengduan | 2014 | muscle | 97.75326 | 24.99136 | 10.5 |
| | DHYJ133 | southern Hengduan | 2014 | muscle | 97.75326 | 24.99136 | 16.1 |
| | DHYJ139 | southern Hengduan | 2014 | muscle | 97.75326 | 24.99136 | 11.5 |
| | DHYJ129* | southern Hengduan | 2014 | muscle | 98.32495 | 27.82627 | 13 |
| *Yuhina diademata* | 2022LS135 | southern Hengduan | 2022 | muscle | 98.713 | 25.95804 | 17.1 |
| | 2022LS151 | southern Hengduan | 2022 | muscle | 98.713 | 25.95805 | 13.3 |
| | 2022LS152 | southern Hengduan | 2022 | muscle | 98.713 | 25.95805 | 11.3 |
| | 2022LS183 | southern Hengduan | 2022 | muscle | 98.68341 | 25.96893 | 15.7 |
| | GLGS2175* | southern Hengduan | 2004 | muscle | 98.65953 | 25.00933 | 12.9 |
| *Yuhina flavicollis* | 2022GS086 | southern Hengduan | 2022 | muscle | 98.35046 | 27.69037 | 15.9 |
| | 2022GS087 | southern Hengduan | 2022 | muscle | 98.35046 | 27.69037 | 12.1 |
| | 2022GS210 | southern Hengduan | 2022 | muscle | 98.35046 | 27.69036 | 13.3 |
| | 2022LS243 | southern Hengduan | 2022 | muscle | 98.71339 | 25.95805 | 13.9 |
| | 2022LS245 | southern Hengduan | 2022 | muscle | 98.71339 | 25.95805 | 17.5 |
| *Yuhina gularis* | GLGS1229 | southern Hengduan | 2003 | muscle | 98.62164 | 25.80947 | 22 |
| | GLGS1248 | southern Hengduan | 2003 | muscle | 98.62133 | 25.81003 | 8.6 |
| | GLGS1398 | southern Hengduan | 2003 | muscle | 98.76033 | 24.85558 | 10.4 |
| | GLGS1399 | southern Hengduan | 2003 | muscle | 98.76033 | 24.85558 | 16.9 |
| | GLGS2174 | southern Hengduan | 2004 | muscle | 98.68167 | 25.97294 | 9.1 |
| *Yuhina occipitalis* | GLGS1218* | southern Hengduan | 2003 | muscle | 98.62164 | 25.80947 | 18 |
| | GLGS1219 | southern Hengduan | 2003 | muscle | 98.62164 | 25.80947 | 9.3 |
| | GLGS1221 | southern Hengduan | 2003 | muscle | 98.62164 | 25.80947 | 10 |
| | GLGS1222 | southern Hengduan | 2003 | muscle | 98.62164 | 25.80947 | 10.1 |
| | GLGS1223 | southern Hengduan | 2003 | muscle | 98.62164 | 25.80947 | 11.6 |
| | 2022LS131 | southern Hengduan | 2022 | muscle | 98.713 | 25.95805 | 18.3 |

| | | | | | | | |
|---|---|---|---|---|---|---|---|
| *Zoothera dixoni* | DHYJ003 | southern Hengduan | 2014 | muscle | 97.64075 | 24.63848 | 16.6 |
| | DHYJ021 | southern Hengduan | 2014 | muscle | 97.62361 | 24.63782 | 15.4 |
| | GLGS5169 | southern Hengduan | 2005 | muscle | 98.70581 | 25.98497 | 15.5 |
| | GLGS1341* | southern Hengduan | 2003 | muscle | 98.67897 | 25.97989 | 20.9 |
| *Zosterops japonicus* | MT460 | eastern Himalayas | 2023 | blood | 95.17747 | 29.24492 | 12.5 |
| | MT464 | eastern Himalayas | 2023 | blood | 95.17747 | 29.24492 | 10.1 |
| | MT467 | eastern Himalayas | 2023 | blood | 95.17747 | 29.24492 | 13.8 |
| | MT469 | eastern Himalayas | 2023 | blood | 95.17747 | 29.24492 | 15.8 |
| | MT672* | eastern Himalayas | 2023 | blood | 98.70486 | 25.98536 | 12.1 |
| *Zosterops palpebrosus* | GLG23150 | southern Hengduan | 2023 | blood | 98.787 | 25.29984 | 10.2 |
| | GLG23185 | southern Hengduan | 2023 | blood | 98.81718 | 25.28924 | 11.2 |
| | GLG23190 | southern Hengduan | 2023 | blood | 98.81654 | 25.28924 | 11.7 |
| | GLG23191 | southern Hengduan | 2023 | blood | 98.80975 | 25.28924 | 14.7 |
| | GLG23195 | southern Hengduan | 2023 | blood | 98.81372 | 25.28924 | 14.5 |
